# Tracking dynamic adjustments to decision making and performance monitoring processes in conflict tasks

**DOI:** 10.1101/2019.12.19.883447

**Authors:** Daniel Feuerriegel, Matthew Jiwa, William F Turner, Milan Andrejević, Robert Hester, Stefan Bode

## Abstract

How we exert control over our decision-making has been investigated using conflict tasks, which involve stimuli containing elements that are either congruent or incongruent. In these tasks, participants adapt their decision-making strategies following exposure to incongruent stimuli. According to conflict monitoring accounts, conflicting stimulus features are detected in medial frontal cortex, and the extent of experienced conflict scales with response time (RT) and frontal theta-band activity in the electroencephalogram (EEG). However, the consequent adjustments to decision processes following response conflict are not well-specified. To characterise these adjustments and their neural implementation we recorded EEG during a modified Flanker task. We traced the time-courses of performance monitoring processes (frontal theta) and multiple processes related to perceptual decision-making. In each trial participants judged which of two overlaid gratings forming a plaid stimulus (termed the S1 target) was of higher contrast. The stimulus was divided into two sections, which each contained higher contrast gratings in either congruent or incongruent directions. Shortly after responding to the S1 target, an additional S2 target was presented, which was always congruent. Our EEG results suggest enhanced sensory evidence representations in visual cortex and reduced evidence accumulation rates for S2 targets following incongruent S1 stimuli. Results of a follow-up behavioural experiment indicated that the accumulation of sensory evidence from the incongruent (i.e. distracting) stimulus element was adjusted following response conflict. Frontal theta amplitudes positively correlated with RT following S1 targets (in line with conflict monitoring accounts). Following S2 targets there was no such correlation, and theta amplitude profiles instead resembled decision evidence accumulation trajectories. Our findings provide novel insights into how cognitive control is implemented following exposure to conflicting information, which is critical for extending conflict monitoring accounts.

## 1. Introduction

In everyday life we are constantly adapting our decision-making strategies to fit the demands of our environment. The cognitive systems that allow us to do this, and their implementation in the brain, have been the focus of extensive investigation. One core area of this research concerns how we deal with conflicting information, specifically in situations where different features of a stimulus are each associated with different motor actions. For example, in the Eriksen Flanker task (Eriksen & Eriksen, 1974) participants must respond based on the direction of a central target arrow, which is flanked by distractor arrows that point in either congruent or incongruent directions to the target. When the target and distractor arrows are incongruent, participants are slower and less accurate in their responding (Eriksen & Eriksen, 1974; Gratton et al., 1992). However, people can rapidly adjust their decision-making strategies to better handle this type of conflict. When the stimulus in the previous trial was incongruent, participants make faster and more accurate responses to incongruent stimuli in the current trial, and are typically slower in responding to congruent stimuli (Gratton et al., 1992; Mayr et al., 2003; Duthoo et al., 2013). This phenomenon is consistently observed across Flanker, Simon and Stroop tasks, and is termed conflict adaptation, the Gratton effect, or congruency sequence effects (Botvinick et al., 2001; Gratton et al., 2017; Schmidt, 2019). Understanding when and how we adjust our decision-making in the face of conflicting information is essential for understanding how we exert control over our behaviour more generally, including in situations where control-related processes lead to impaired performance (e.g., Wessel, 2017).

A class of highly influential ‘conflict monitoring’ models (Botvinick et al., 2001, 2004; Shenhav et al., 2013) have been developed to account for congruency sequence effects. According to these models, changing task demands are detected and signalled by structures in medial prefrontal cortex based on the presence of conflict. This conflict signalling is indexed by increased fMRI BOLD signals in the anterior cingulate cortex (ACC, reviewed in Shenhav et al., 2013) and increases in theta-band (4-8 Hz) spectral activity in electrophysiological recordings (e.g., Cohen & Donner, 2013; reviewed in Cohen, 2014). In Flanker tasks, conflict is defined as co-activation of neural populations in motor areas that correspond to incompatible motor actions (Cohen, 2014; but see Brown & Braver, 2008 for an alternative definition). Strategic adjustments to decision-making processes are then implemented by a distributed network across prefrontal and parietal areas (Cavanaugh & Frank, 2014). The adjustments described in these models relating to congruency sequence effects are trial-by-trial shifts in the attention allocated to different stimulus features, depending on the expected utility of each stimulus feature for performing the task at hand (Botvinick et al., 2001; Gratton et al., 2017). For example, in trials where distractors cue the incorrect choice, attention to these distractors is reduced in the following trial, which in turn also reduces the detrimental effects of target/distractor incongruence on accuracy and response times (RTs). A second consequence of this adjustment is that participants do not as effectively take advantage of the information conveyed by distractors in trials where they are congruent with the target and cue the correct response (Gratton et al., 1992; Shenhav et al., 2013).

How this theorised attention shifting mechanism operates, and how it influences those processes which are active during perceptual decision-making, are not well specified in these models. It is unclear whether attention refers to spatial or feature-based attention, corresponding to response gain changes in stimulus-selective sensory neurons (e.g., Reynolds & Heeger, 2009), or changes to how information provided by sensory cortex is used to make a decision, which may instead occur across a network of parietal and prefrontal areas (e.g., Afacan-Seref et al., 2018). It is also unclear whether other adjustments associated with cognitive control are implemented following response conflict, such as changes in the amount of sensory evidence required to make a decision (e.g., Forstmann et al., 2008). Without extending conflict monitoring models to describe these adjustments it is difficult to meaningfully test them against competing accounts, such as those which explain congruency sequence effects as driven by associative learning of stimulus-response associations (Hommel et al., 2004; Abrahamse et al., 2016; Schmidt, 2019). Accordingly, the primary aim of the current study was to develop a framework for tracking rapid adjustments to decision-making processes that occur following response conflict, in order to develop more specific and testable versions of conflict monitoring models.

To characterise the adjustments that occur following response conflict, we recorded electroencephalography (EEG) while participants completed a novel variant of the Flanker task (here termed a modified Flanker task). In each trial participants responded to two target stimuli, termed the S1 and S2 targets. The first target could be either a congruent or incongruent stimulus. Shortly after responding to this S1 target, the S2 target was presented, which was always congruent. Importantly, the correct responses corresponding to the S1 and S2 targets were not dependent on each other (i.e. participants made two independent perceptual decisions in each trial). Here, we assessed whether patterns of behavioural and neural responses for S2 targets differed by S1 congruency (note that ‘congruency’ in our study refers to the target/distractor elements within S1 and S2, respectively, and not the relation between S1 and S2). We adopted an EEG analysis framework (developed by O’Connell et al., 2012; Steinemann et al., 2018) that allowed us to simultaneously track the neural correlates of multiple processes that are critical for perceptual decision-making. This approach is directly inspired by evidence accumulation models of decision-making, such as the diffusion model (Ratcliff, 1978; Ratcliff & Smith, 2004; Ratcliff et al., 2016), which formalise perceptual decision-making as a process whereby sensory evidence in favour of each decision outcome is accumulated over time. When this evidence reaches a set threshold, or ‘decision bound’, associated with a decision outcome, the motor action corresponding to that decision is initiated. Using our EEG analysis framework, we could trace the time-courses of multiple processes described in these models, including the representation of sensory evidence in visual cortex, the build-up of decision evidence, and preparatory motor activity corresponding to different decision alternatives. Critically, the adjustments to decision-making processes that occur following response conflict should be indexed by effects on these EEG measures. Our approach also facilitates identification of distinct adjustments at different stages of the decision-making process, each of which may have opposing effects on accuracy and response speed, which are difficult to identify based on behavioural data alone (see Steinemann et al., 2018; Kelly et al., 2020).

To track the representation of decision-relevant sensory evidence we presented stimuli that consisted of overlaid left and right tilted gratings and asked participants to judge which grating was of dominant (i.e. of higher contrast, as done by Steinemann et al., 2018). In our modified Flanker task we presented S1 target and distractor stimuli that could have dominant gratings in congruent or incongruent directions. The distractor consisted of a central annulus that encircled a fixation cross, and the target consisted of a larger annulus that encircled the distractor. This resulted in a difficult Flanker task that required participants to ignore the centrally-presented distractor in incongruent trials and make decisions based on the more peripheral target. In contrast to more typical Flanker tasks, the distracting information was presented near fixation, and the target stimulus was presented in the periphery. Each grating contrast reversed at different periodicities (at 15 Hz and 20 Hz), allowing us to separately tag visual responses to each grating by measuring steady-state visual evoked potentials (SSVEPs; Regan, 1966; Norcia et al., 2015). Importantly, SSVEP amplitudes monotonically scale with the contrast of each grating (Campbell & Kulikowski, 1972; Allen et al., 1986) and are modulated by spatial and feature-based attention (Morgan et al., 1996; Muller et al., 2006). As the contrast levels of each grating were the critical stimulus features in our task, SSVEP magnitudes corresponding to each grating orientation were closely correlated with the extent of sensory evidence for each decision outcome. Accordingly, we calculated measures of sensory evidence favouring the correct response, defined as the SSVEP amplitude favouring the stronger target grating orientation (associated with the correct response) minus that evoked by the weaker grating orientation. This captured the sum of sensory evidence favouring the correct response across the target and distractor elements of the stimulus (for a similar approach see Steinemann et al., 2018).

We also tracked the accumulation of decision evidence over time by measuring the build-up rate of the centro-parietal positivity (CPP) event-related potential (ERP) component (O’Connell et al., 2012). The shape of this component corresponds closely with the trajectory of decision evidence accumulation in diffusion models (Twomey et al., 2015; Kelly et al., 2020). In addition, we measured spectral amplitude changes in the Mu/Beta (8-30 Hz) range at electrodes over left and right motor cortex, which index the build-up of preparatory motor activity preceding a behavioural response, such as a keypress (Donner et al., 2009; de Lange et al., 2013). This motor preparation response typically displays similar accumulation-to-bound dynamics as the CPP and reaches a set threshold just before the execution of a motor action (e.g., O’Connell et al., 2012). However, when there are strict response deadlines that induce speed pressure (as commonly used in conflict tasks, e.g., Cohen & Donner, 2013; Tollner et al., 2017) motor preparation can begin ramping toward the motor action threshold even before a target stimulus is presented, and continues to ramp toward this threshold until a motor action is executed (Steinemann et al., 2018; Kelly et al., 2020). The level of motor activity determines when and if a motor action is made, and so this ramping motor activity can produce behavioural responses that are based on less decision evidence than if there was no speed pressure (Steinemann et al., 2018; Kelly et al., 2020). In these contexts, adjustments to pre-target levels of motor activity can produce decision-making strategies that favour either fast or accurate responses.

By simultaneously tracking each of these neural markers, we could identify the adjustments to decision-making strategies that correspond to attention shifts as described in conflict monitoring models. If these attention shifts are associated with spatial or feature-based attention in visual cortex (whereby response gain is reduced specifically for neurons responsive to the distractor) then we would expect to observe smaller SSVEP sensory evidence measures evoked by the (always congruent) S2 targets when the previous stimulus was incongruent, accompanied by a shallower build-up rate of the CPP. If the attention shift relates to how sensory information is utilised after it is transmitted from visual cortex, then we would expect to observe slower CPP build-up rates following incongruent stimuli, but without co-occurring reductions in SSVEP amplitudes. If there are also shifts in pre-target motor response preparation following response conflict, then we would expect to see shifts in pre-target Mu/Beta amplitudes. Based on the notion that attention shifts in conflict monitoring accounts reflect changes in spatial attention (e.g., Janssens et al., 2017), we expected to observe slower RTs to S2 targets following incongruent S1 targets (a post-conflict slowing effect for congruent stimuli; Gratton et al., 1992; Mayr et al., 2003), accompanied by smaller SSVEP measures of sensory evidence (corresponding to reduced attention to the central distractor stimulus) and slower rates of decision evidence accumulation as tracked by the CPP.

We further extended this analysis framework to measure theta-band (4-8 Hz) activity at midfrontal electrode FCz. Larger frontal theta amplitudes for incongruent stimuli in the ∼400 ms time window immediately preceding a behavioural response have been consistently observed across experiments (e.g., Cohen & Cavanaugh, 2011; Cohen & Donner, 2013; van Driel et al., 2015). We included this measure to assess whether similar markers of conflict signalling could be found when using our modified Flanker task, and also to see whether frontal theta amplitudes for S2 targets would be modulated by conflict at S1 (as reported by Jiang et al., 2018).

This also allowed us to assess the temporal profiles of frontal theta amplitudes in relation to other neural correlates of decision-making processes. Notably, frontal theta activity has been proposed as a neural correlate of decision evidence accumulation, as it shows similar accumulation-to-bound profiles to the CPP (van Vugt et al., 2012). However, frontal theta follows a very different temporal profile in conflict tasks (Cohen & Cavanaugh, 2011; Cohen & Donner, 2013; van Driel et al., 2015; Tollner et al., 2017) and in some perceptual decision tasks that do not involve response conflict (Werkle-Bergner et al., 2014), whereby frontal theta steadily rises in amplitude at a fixed rate from target onset until the time of the behavioural response. By simultaneously tracking multiple neural markers of decision processes, we observed two potential contributions to frontal theta power in addition to effects that are specifically associated with the detection of conflict.

To foreshadow our results, we identified reductions in the rate of evidence accumulation following response conflict, indexed by build-up rates of the CPP following S2 targets. Further evidence congruent with changes in evidence accumulation rates was observed in a follow-up behavioural experiment. These findings provide initial evidence of how cognitive control is implemented across decision-making circuits following response conflict.

## 2. Methods

### 2.1 Participants

35 people (24 female, 11 male), aged between 18-40 (M = 24.1) took part in this experiment. Participants were right-handed, fluent in English and had normal or corrected-to-normal vision. 6 participants were excluded from analyses because they achieved less than 60% accuracy in at least one experimental condition. An additional 2 participants were excluded from the EEG dataset: one due to excessively noisy data, and one due to data loss following a technical error. This resulted in a sample of 29 participants for behavioural data analyses, aged between 18-36 (M = 23.9), and 27 for EEG analyses. Participants were compensated 25 AUD for their time. This study was approved by the Human Ethics Committee of the Melbourne School of Psychological Sciences (ID 1750871).

### 2.2 Stimuli

Stimuli were presented using a gamma-corrected 24” LCD monitor with a refresh rate of 60 Hz. Stimuli were presented using functions from MATLAB (Mathworks) and PsychToolbox (Brainard, 1997; Kleiner et al., 2007). Code used for stimulus presentation will be available at https://osf.io/eucqf/ at the time of publication.

The critical stimuli consisted of two overlaid gratings within a circular aperture, presented against a grey background, similar to stimuli in Steinemann et al. (2018). Each of the two gratings were oriented 45° to the left and right of vertical, respectively (Figure 1B). The left-tilted grating contrast-reversed at a rate of 15 Hz; the right-tilted grating contrast-reversed at a rate of 20 Hz. The circular aperture was divided into two concentric circles; an inner circle and an outer circle with radii of 1.32° and 4.70° of visual angle, separated by a grey spacer ring.

**Figure 1.**
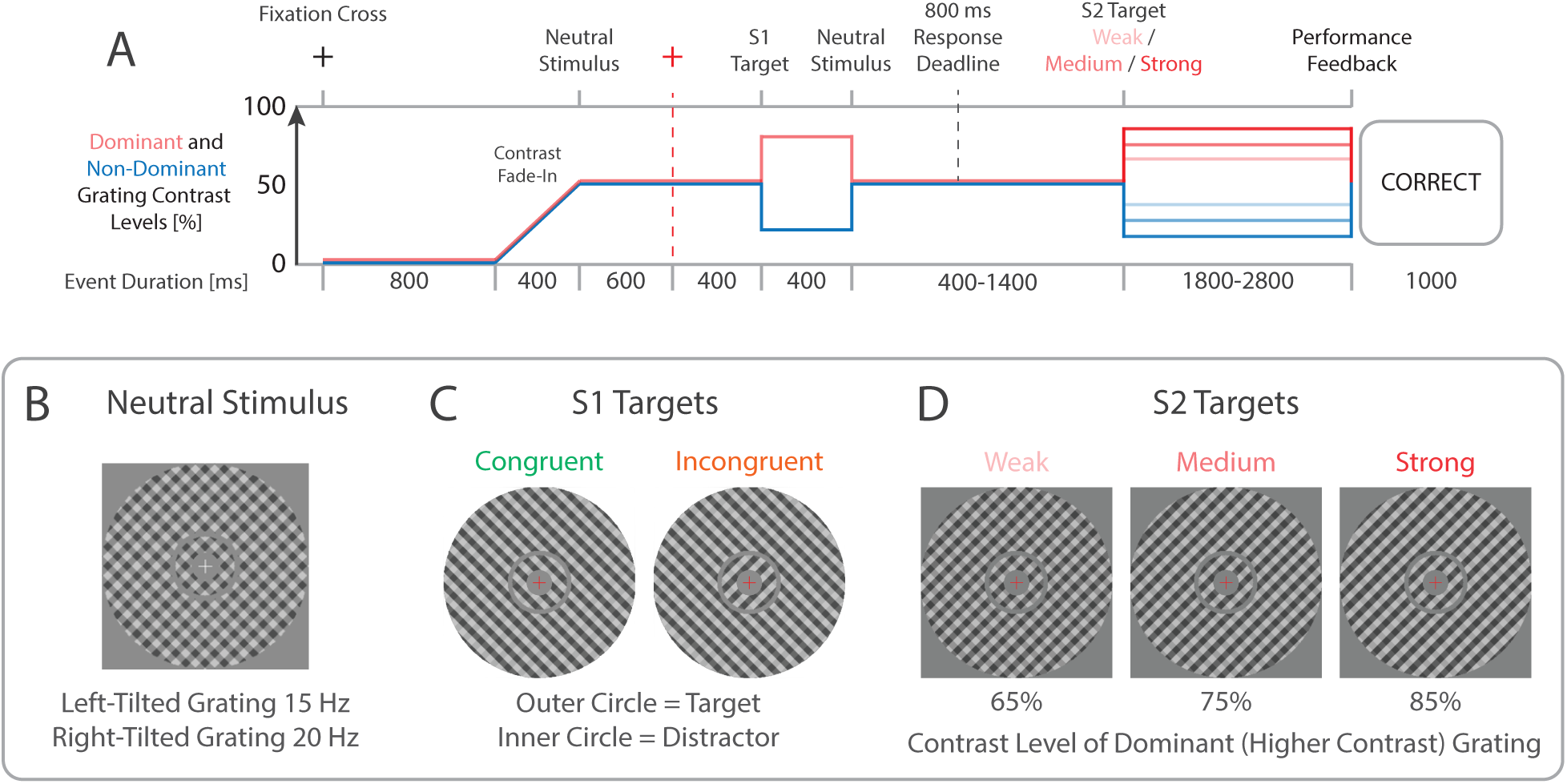
Trial structure and task. A) Each trial commenced with the presentation of a fixation cross. Following this, two circular apertures containing overlaid left- and right-tilted gratings were presented. Red and blue lines respectively depict the contrast levels of the dominant (i.e. higher contrast) and non-dominant (lower contrast) gratings in each phase of the trial. B) The left- and right-tilted gratings contrast reversed at 15 Hz and 20 Hz, respectively, and were each presented at 50% contrast (Neutral Stimulus). The fixation cross then changed to red, and 400 ms later the relative contrast levels of the two gratings were changed to create the S1 target. C) The outer circle was the target stimulus, and the inner circle was the distractor. Participants indicated which set of stripes in the target stimulus was dominant (i.e. of higher contrast). In trials with congruent distractors the dominant grating orientations were the same for the inner and outer circles. In trials with incongruent distractors they were of opposite orientations. Participants were required to respond within 800 ms following S1 target onset, after which a neutral stimulus was presented for a further 600-800 ms. D) Following this the S2 target appeared, which always consisted of a target and congruent distractor, and was presented until the response deadline at the end of the trial. The direction of the dominant grating in the S2 target was not dependent on that of the S1 target. The contrast level of the dominant grating could be either weak (65%) medium (75%) or strong (85%) for both target and distractor elements. After each trial participants received feedback on their response to the S2 target, or feedback indicating they had responded too early or missed the S1 response deadline.

### 2.3 Procedure

Participants sat 100 cm from the monitor in a darkened room and were asked to fixate on a central cross throughout all trials. Each trial included two task phases; each task phase comprised a single target that required a response. The trial structure is depicted in Figure 1A. In each trial, a white fixation cross appeared for 800 ms. Following this, both gratings gradually increased in contrast from 0% to 50% over a 400 ms period. Both gratings remained at 50% contrast for a further 1000 ms, during which the contrast levels of both gratings were identical (i.e. the stimulus was “neutral”, see Figure 1B). 400 ms before the end of this period the central fixation cross changed colour to red, signifying that the first target (here termed the S1 target) would soon appear. Immediately after this neutral stimulus period, one of the gratings within each circle increased to 80% contrast and the other decreased to 20% contrast. This contrast difference persisted for 400 ms, after which the neutral stimulus was presented. In congruent trials, the stripes of higher contrast were the same orientation for inner and outer circles; in incongruent trials they were of opposite orientations (Figure 1C). Congruent and incongruent trials were each presented with 50% probability. Participants indicated which grating was dominant (i.e. of higher contrast) in the outer circle (while ignoring the inner circle) by pressing keys on a TESORO Tizona Numpad (1000 Hz polling rate) using their left and right index fingers. Participants were required to respond within 800 ms of S1 target onset. Following the response to S1 the neutral stimulus was presented for a further 600 ms.

Following this neutral stimulus period, the second (S2) target appeared at the time of the next screen refresh when the phase-reversals of both gratings were in synchrony, which occurred every 12 frames [200 ms]. The S2 target always consisted of targets and congruent distractors, however the contrast difference between the higher and lower contrast gratings varied across 3 levels: weak (one grating 65% contrast, the other 35%), medium (75/25%) and strong (85/15%; for examples see Figure 1D). Participants indicated via keypress which grating was dominant, as done following the S1 target. The S2 target was presented for durations ranging from 1800-2400 ms depending on the timing of the S1 response, so that the total presentation duration of the gratings (including both S1/S2 target and neutral stimulus periods) was equated across trials. Importantly, the direction of the dominant grating in the S2 target was independent of the direction of the S1 target within the same trial, and participants were explicitly notified of this.

A feedback screen was then displayed for 1000 ms. Participants received “Correct” or “Error” feedback depending on their response to the S2 target. If responses were made prior to the S1 target or after the 800 ms S1 response deadline, then “Too Early” of “Too Slow” feedback appeared instead. If no response was given following the S2 target, then “No Response” feedback was presented. Correct/error feedback relating to the S1 target was not provided, so that participants would not keep their response to the S1 target in memory and match this to the feedback presented, which may have interfered with subsequent decisions related to the S2 targets.

Participants completed 480 trials, split into 10 blocks of 48 trials each. Participants were allowed self-paced breaks between blocks (minimum break duration 15 seconds). Prior to the experimental blocks, participants completed a practice block of 12 trials, which was repeated until participants demonstrated adequate performance. During this block, participants received feedback on their responses to both S1 and S2 targets in each trial. Trial order was randomised and proportions of trials with each S1 and S2 target type combination were balanced within each block.

### 2.4 Analyses of Accuracy and RT Data

Trials with responses that were too slow or earlier than stimulus onset were removed from the dataset. Only trials with correct responses and RTs of >100 ms were included for analyses of RTs. We modelled proportions of correct responses using generalised linear mixed effects logistic regressions (binomial family) as implemented in the R package lme4 (Bates et al., 2015). We modelled RTs using generalised linear mixed effects regressions using a Gamma family and an identity link function, as recommended by Lo & Andrews (2015). Given that error RTs are also diagnostic of decision processes within evidence accumulation model frameworks (e.g., Ratcliff & Smith, 2004), we have also plotted mean RTs for errors in Supplementary Figure S1.

To test for effects of each factor of interest on accuracy and RT measures, we compared models with and without that fixed effect of interest using likelihood ratio tests. For each comparison, both models included all other fixed effects that would conceivably influence the data, as well as identical random effects structures. Fixed effects of interest for S1 targets included S1 congruency (congruent, incongruent). In comparison models we also included effects of S1 target orientation (left, right). Fixed effects of interest for S2 targets included S1 congruency (congruent, incongruent) and S2 evidence strength (strong, medium, weak). Additional effects included in both models with and without each fixed effect of interest included additive and interactive effects of S1 and S2 target orientation (left, right). The structure of each model and the coefficients of each fitted model are detailed in the Supplementary Material.

### 2.5 EEG Data Acquisition and Processing

We recorded EEG at a sampling rate of 512 Hz from 64 active electrodes using a Biosemi Active Two system (Biosemi). Recordings were grounded using common mode sense and driven right leg electrodes (http://www.biosemi.com/faq/cms&drl.htm). We added 6 additional channels: two electrodes placed 1 cm from the outer canthi of each eye, and electrodes placed above and below the center of each eye.

We processed EEG data using EEGLab v13.4.4b (Delorme & Makeig, 2004). All data processing and analysis code and corresponding data will be available at https://osf.io/eucqf/ at the time of publication. First, we identified excessively noisy channels by visual inspection (median number of bad channels = 1, range 0-7) and excluded these from average reference calculations and Independent Components Analysis (ICA). Sections with large artefacts were also manually identified and removed. We re-referenced the data to the average of all channels, low-pass filtered the data at 30 Hz (EEGLab Basic Finite Impulse Response Filter New, default settings), and removed one extra channel (AFz) to correct for the rank deficiency caused by the average reference. We processed a copy of this dataset in the same way and additionally applied a 1 Hz high-pass filter (EEGLab Basic FIR Filter New, default settings) to improve stationarity for the ICA.

ICA was performed on the high-pass filtered dataset (RunICA extended algorithm, Jung et al., 2000). We then copied the independent component information to the unfiltered dataset (e.g., as done by Feuerriegel et al., 2018). Independent components generated by blinks and saccades were identified and removed according to guidelines in Chaumon et al. (2015). After ICA we interpolated any excessively noisy channels and AFz using the cleaned data (spherical spline interpolation). EEG data were then high-pass filtered at 0.1 Hz (EEGLab Basic Finite Impulse Response Filter New, default settings).

The resulting data were segmented from −2200 ms to 3800 ms relative to the S1 target onset, and baseline-corrected using the interval of 400-600 ms prior to the S1 target. This baseline period was used to ensure that ERPs evoked by the fixation cross colour change did not influence baseline estimates. Epochs containing amplitudes exceeding ±150 μV at any scalp channels were rejected (mean trials retained = 462 out of 480, range 413-479). Numbers of retained epochs by condition are displayed in Supplementary Table S1. Data were then converted to current source density (CSD) estimates using the CSD Toolbox (Kayser & Tenke, 2006; m-constant = 4, λ = 0.00001). The resulting S1 target-locked epochs were used for all subsequent EEG analyses. Only data from trials with correct responses to S1 targets were used for analyses of S1 neural responses, as is typical in EEG analyses of conflict task data (e.g., Cohen & Donner, 2013). Similarly, only trials with correct responses to both S1 and S2 targets were included in analyses of S2 neural responses, to exclude effects of errors and post-error adaptations (Wessel, 2017).

We then segmented the EEG data according to four time windows of interest: from −500 ms to 1000 ms relative to S1 target onset, from −1000 ms to 500 ms relative to S1 responses, from −200 ms to 1000 ms relative to S2 target onset, and from −700 ms to 300 ms relative to S2 responses. Data were transformed into frequency domain representations (using data from the entire trial) before these epochs were derived for SSVEP, Theta and Mu/Beta analyses.

An overview of the EEG data analysis approach is depicted in Figure 2. Each EEG measure (SSVEPs, CPP, mu/beta and frontal theta-band activity) corresponded to successive stages of the hypothesised decision-making process (shown in Figure 2A). These measured were compared across S1 congruency and S1 RT quantile conditions following S1 targets, and across S1 congruency and S2 evidence strength conditions following S2 targets (Figure 2B).

**Figure 2.**
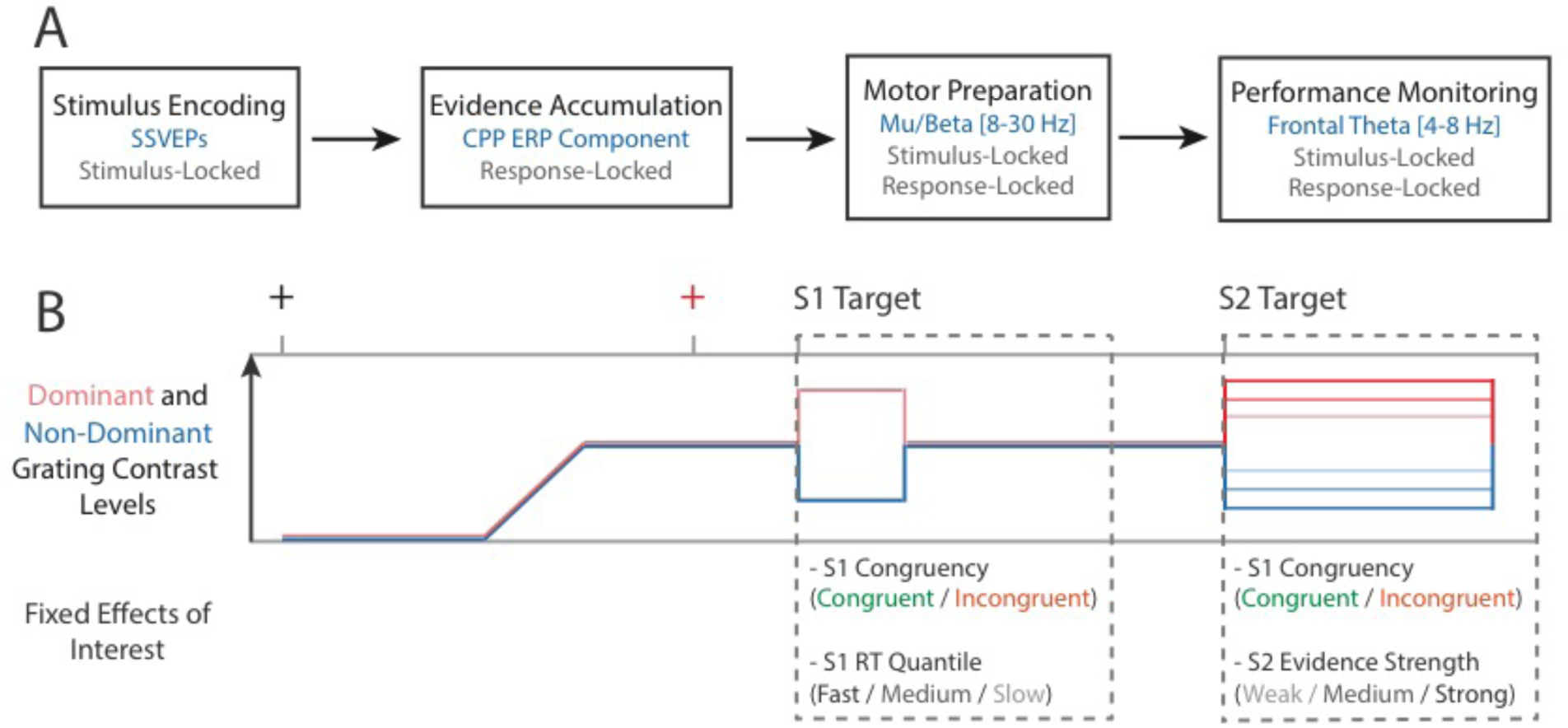
Overview of the EEG data analysis approach. A) Neural correlates of successive stages of the decision-making process were measured. These stages included stimulus encoding (indexed by SSVEPs), evidence accumulation (indexed by the CPP ERP component), motor action preparation (indexed by mu/beta activity over motor cortex) and performance monitoring or conflict detection (indexed by frontal theta-band activity). Grey text denotes whether each measure was measured relative to stimulus onset, the time of the response, or relative to both of these time points. B) Different fixed effects of interest were tested in the time windows following S1 and S2 targets. For the time window following S1 targets effects of S1 congruency and S1 RT quantile were of interest. For the time window following S2 targets effects of S1 congruency and S2 evidence strength were of interest.

### 2.6 Analyses of SSVEPs

SSVEP amplitudes were estimated using data from electrode Oz. We converted data into the frequency domain using Fast Fourier transforms (FFTs; window width 400 ms, step size 10 ms, Hann taper). Amplitudes at frequency bins corresponding to the 15 Hz and 20 Hz contrast reversal rates were baseline-corrected by subtracting the average amplitude of the two neighbouring frequency bins (see Norcia et al., 2015; Retter & Rossion, 2016). Then, a measure of evidence in favour of the dominant target grating orientation was derived by subtracting the SSVEP signal corresponding to the higher contrast grating from that of the lower contrast grating. Importantly, both the target and distractor elements of the stimulus contributed to the SSVEPs corresponding to each grating orientation, and so these measures index the sum of evidence across target and distractor elements. To avoid biases due to differences in trial numbers across targets with different flicker frequencies and S1/S2 orientation conditions, we averaged signals across trials of each S1 and S2 target orientation combination separately, and then averaged the resulting signals within each condition of interest.

For S1 target-evoked SSVEPs we compared congruent and incongruent conditions. For S2 target-evoked SSVEPs we tested for differences by S1 congruency and S2 evidence strength (weak/medium/strong). Differences in the magnitude of sensory evidence favouring the correct response across conditions were tested for using mass-univariate analyses. Paired-samples t tests were performed at all time steps between −500 ms and 1000 ms relative to the S1 target, and between −200 ms and 1000 ms relative to the S2 target. To correct for multiple comparison, cluster-based permutation tests using the cluster mass statistic (Bullmore et al., 1999; Maris and Oostenveld, 2007; 1000 permutation samples, cluster formation alpha = 0.05) were performed using functions from the Decision Decoding Toolbox v1.0.3 (Bode et al., 2019).

### 2.7 Analyses of ERPs

We measured the slopes and pre-response amplitudes of the CPP component at electrode Pz. To measure CPP slopes we first averaged ERPs across response-locked epochs within each condition. We then fit a regression line to data ranging from −350 ms to −50 ms relative to the time of the keypress response. This time window was selected to capture evidence accumulation dynamics during the vast majority of trials (as done by Steineimann et al., 2018). CPP slopes were compared across conditions using paired-samples t tests and repeated measures ANOVAs implemented in JASP v0.9.1 (JASP Core Team). Pre-response CPP amplitudes were measured as the average amplitude between −130 ms and −70 ms from response onset. This time window was chosen to capture the amplitude of the CPP around the onset time of motor execution prior to completion of the keypress (as defined in Steinemann et al., 2018). We analysed CPP pre-response amplitudes using linear mixed effects regression models and tested for effects using likelihood ratio tests as described above. Fixed effects of interest for S1 targets included S1 congruency and S1 RT. Fixed effects of interest for S2 targets included S1 congruency and S2 evidence strength. Additional effects in both models with and without each fixed effect of interest included additive and interactive effects of S1 and S2 target orientation. The structures of all models used in these analyses are detailed in the Supplementary Material.

### 2.8 Analyses of Frontal Theta and Mu/Beta Amplitudes

Time-frequency amplitude estimates were derived using complex Morlet wavelet convolution, using a frequency range of 1-30 Hz, and linear steps of 1 Hz. The number of wavelet cycles increased linearly from 3-10 cycles across this frequency range. Time-frequency amplitude estimates were converted to decibels (dB) relative to the median of amplitudes across all trials with correct S1 responses for each participant, averaged across a baseline period from −900 ms to −700 ms from S1 target onset. This baseline period was used to minimise contributions of the fixation cross colour change to baseline estimates. S2 stimulus-locked epochs and S1 and S2 response-locked epochs were created using this decibel-transformed data.

Frontal theta responses were measured as the average of dB estimates across 4-8 Hz at electrode FCz (Cohen & van Gaal, 2014). Mu/Beta responses associated with motor preparation were measured as the average across 8-30 Hz at electrodes C3 and C4 (Steinemann et al., 2018). Motor preparation signals at electrodes that were contralateral and ipsilateral to the response hand used in each trial were analysed separately, as these signals show different dynamics preceding an executed motor response (Donner et al., 2009; de Lange et al., 2013).

Frontal theta and Mu/Beta amplitudes were averaged across trials for each S1 and S2 target orientation combination separately, and then averaged across all S1/S2 combinations within each condition of interest. We performed mass-univariate paired-samples t-tests and corrections for multiple comparisons using cluster-based permutation tests as described above. We did not have specific *a priori* hypotheses about how Frontal theta or Mu/Beta responses would vary by S2 target evidence strength, and instead performed post-hoc mass-univariate comparisons to test for differences between strong and weak evidence conditions.

## 3. Results

### 3.1 Behavioural Results

We observed typical effects of response conflict on behaviour. Participants were slower and less accurate in responding to S1 targets with incongruent distractors (likelihood ratio test p’s < 0.001, Figures 3A, 3B). The delta plot in Figure 3C displays larger effects of congruency on RTs for slower RT quantiles. This pattern is typical of both Flanker and Stroop tasks (see Ulrich et al., 2015), but differs from delta functions observed in some variants of Simon tasks (e.g., Ridderinkhof, 2002), which show smaller delta values for slower RT quantiles (reviewed in Pratte et al., 2010; Proctor et al., 2011; Ulrich et al., 2015). Consistent with task conditions involving strict response deadlines, RTs for error trials appeared to be slightly faster than correct responses, particularly in trials with congruent stimuli (depicted in Supplementary Figure S1).

**Figure 3.**
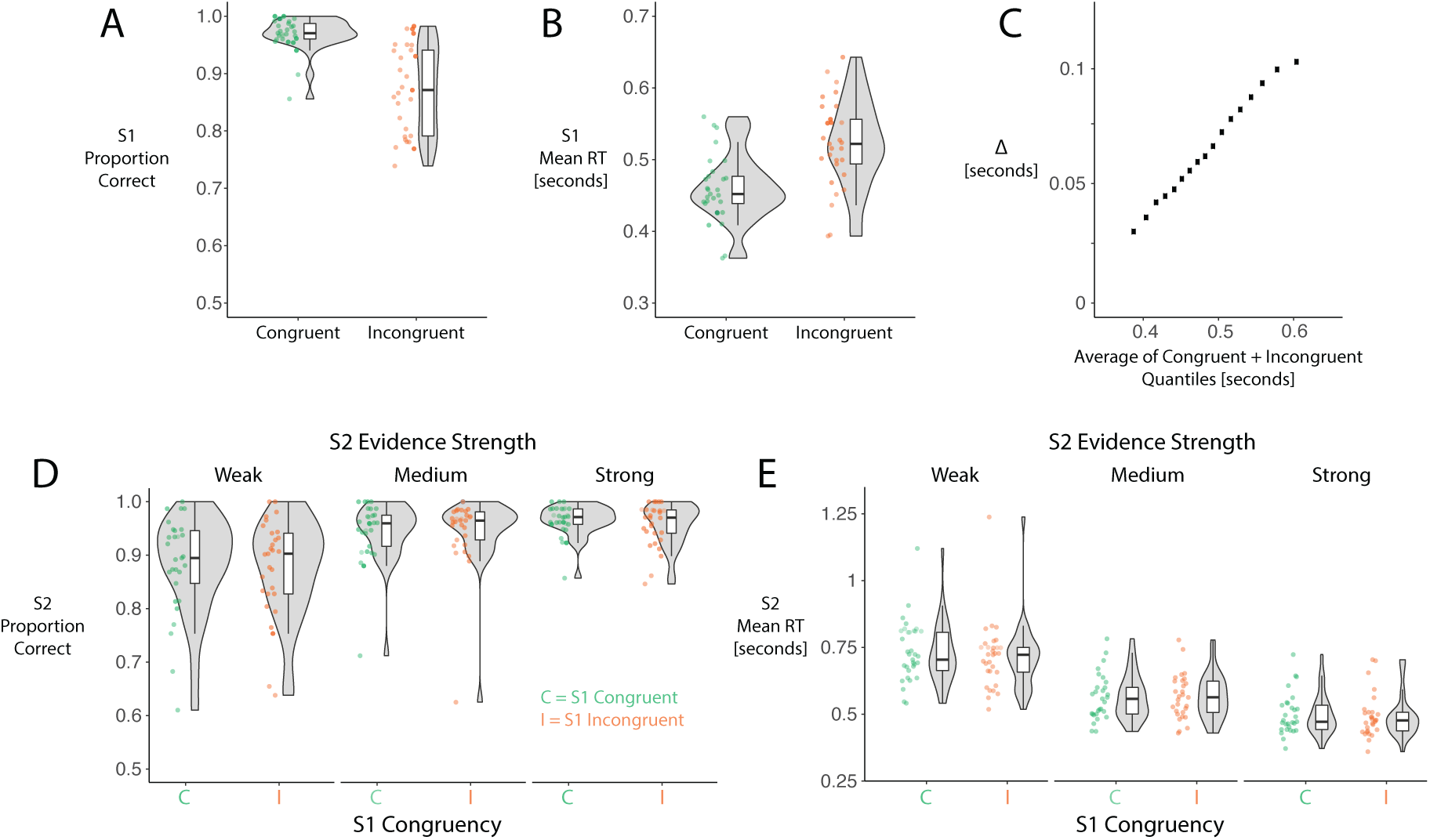
Accuracy and mean RTs following S1 and S2 targets. A) Proportions of correct responses following S1 targets with congruent and incongruent distractors. B) Mean RTs for correct responses to S1 targets with congruent and incongruent distractors. C) Delta plot displaying the [incongruent – congruent] differences in RTs for 10-90% quantiles in steps of 5% on the Y axis and the average of each congruent and incongruent condition RT quantile on the X axis. D) Proportions of correct responses to S2 targets, split by S2 evidence strength and S1 distractor congruency. E) Mean RTs for correct responses to S2 targets.

Following S2 targets, participants were faster and more accurate in trials with targets of higher evidence strength (p’s < 0.001, Figures 3D, 3E). However, the addition of S1 congruency did not significantly improve model fits for accuracy (p = 0.484) or RT data (p = 0.099). The estimate of the S1 distractor congruency effect indicated a tendency toward post-conflict speeding rather than slowing (fixed effect of congruency = −6.2 ms).

### 3.2 Neural Responses Following S1 Targets

Effects of S1 congruency were clearly visible in all EEG measures, except for motor preparation-related Mu/Beta amplitudes. These effects are depicted in Figure 4, and are consistent with previous findings (e.g., Cohen & Donner, 2013; Jiang et al., 2018). SSVEP-based estimates of sensory evidence magnitudes favouring the correct response were smaller following S1 targets with incongruent distractors from ∼50-500 ms after target onset (Figure 4A). This is to be expected, given that the distractor contained higher contrast gratings of the opposite orientation to the target, and both the target and distractor contribute to the SSVEP measures. Estimates of sensory evidence magnitudes were also smaller in trials with slower RTs, as visible for both congruent and incongruent conditions (Figure 4A). To verify this, we tested whether SSVEP measures in trials with RTs in the fastest tertiles were larger than those in the slowest tertiles, using a one-tailed cluster-based permutation test. Here, we took SSVEP measures from trials with RTs in the fastest/slowest RT tertiles, for congruent and incongruent distractor conditions separately, and then averaged SSVEP measures across congruency conditions within each participant. We found a significant cluster ranging from ∼0-400 ms following S1 target onset. Note that this test was done after observing the data, and such tests are circular and have an inflated false positive rate (see Kriegeskorte et al., 2009), but that our results are similar to previously-observed patterns in Steinemann et al. (2018).

**Figure 4.**
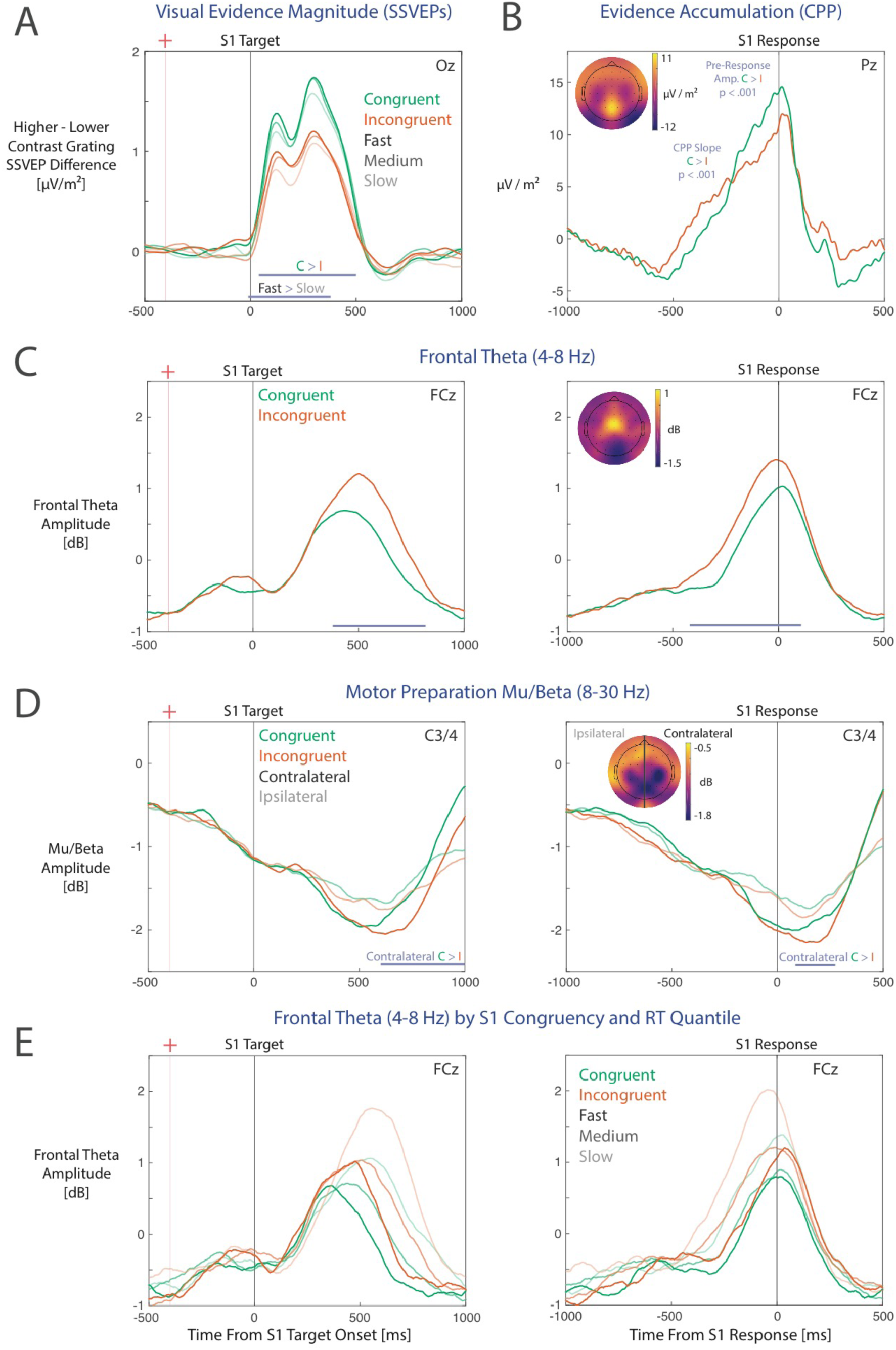
Neural responses following S1 targets with congruent and incongruent distractors. A) SSVEP difference measures indexing the magnitude of sensory evidence in favour of the dominant (i.e. higher contrast) target grating, also plotted by fast/medium/slow RT quantiles. Purple bars denote clusters of time points at which there were statistically significant differences across conditions. B) CPP amplitudes leading up to the time of the keypress response. C) Frontal theta amplitudes relative to the target onset (left panel) and response times (right panel). D) Mu/Beta amplitudes indexing motor preparation for electrodes contralateral and ipsilateral to the response hand. E) Frontal theta amplitudes plotted by congruency and RT quantile. Scalp maps depict the average magnitudes of each signal over the 200 ms period preceding the keypress response, averaged across S1 congruent and incongruent conditions.

As expected based on the patterns of SSVEP results, CPP slopes preceding responses to incongruent stimuli were shallower, t(26) = 6.81, p < 0.001. Additionally, pre-response CPP amplitudes were smaller in trials with incongruent stimuli (likelihood ratio test p < 0.001). Pre-response CPP amplitudes were also smaller in trials with longer RTs (p < 0.001). We also performed post-hoc comparisons of CPP slopes from trials with RTs in the fastest and slowest RT tertiles, and found that the build-up rate of the CPP was steeper in trials with faster RTs, t(26) = 4.62, p < 0.001 (see also O’Connell et al., 2012; Twomey et al., 2015).

Frontal theta-band (4-8 Hz) amplitudes were larger following incongruent stimuli from 400-800 ms relative to target onset, and from −450 to 50 ms relative to the time of the keypress response (Figure 4C; for time-frequency plots see Supplementary Figure S2), similar to that reported in previous studies using conflict tasks (e.g., Cohen & Donner, 2013).

Plotting frontal theta amplitudes by S1 distractor congruency and fast/medium/slow RT quantiles (displayed in Figure 4E) allowed us to better characterise the temporal dynamics of this response (as done by Murphy et al., 2015). Theta amplitudes gradually increased at a fixed rate over the course of the trial until ∼100 ms before the time of the response, after which frontal theta amplitudes rapidly decreased (see also Figure 10 in Cohen & Cavanaugh, 2011; Figure 2 in van Driel et al., 2015). This build-up was terminated earlier in trials with faster RTs, leading to lower pre-response theta following congruent S1 targets, as participants responded more quickly in this condition (see also Figure 6 in Tollner et al., 2017).

Motor preparation activity had a different temporal profile to the CPP, showing a steady increase in activity (i.e. more negative amplitudes over time) at electrodes both contralateral and ipsilateral to the response hand. This occurred from the time of the fixation cross colour change preceding the target, which resulted in pre-response amplitudes almost halfway from zero dB to the −2dB motor execution threshold by the time of the S1 target onset (see also Steinemann et al., 2018; Kelly et al., 2020). Mu/Beta amplitudes did not appear to diverge between contralateral and ipsilateral electrodes until around 200 preceding the time of the response (contralateral minus ipsilateral measures plotted in Supplementary Figure S3). However, there were no significant effects of S1 congruency on motor preparation activity until 50-250 ms after the keypress response (Figure 4D). In addition, motor activity at electrodes contralateral to the response hand reached the same threshold at the time of the response for both congruent and incongruent conditions (O’Connell et al., 2012; Kelly et al., 2020).

### 3.3 Neural Responses Following S2 Targets

#### 3.3.1 Effects of S1 congruency

Although we did not find group-level differences in accuracy or RTs to S2 targets by S1 congruency, there was evidence of two opposing adjustments at different stages of the decision process. Sensory evidence magnitudes in favour of the correct response (as indexed by SSVEPs) were larger following S2 targets that appeared after incongruent S1 stimuli between ∼200-400 ms post S2 target onset (Figure 5A). However, CPP build-up rates were slightly slower following incongruent S1 stimuli, t(26) = 2.25, p = 0.033 (Figure 5B) indicative of a slower rate of decision evidence accumulation (Twomey et al., 2015). Pre-response CPP amplitudes did not appear to differ between S1 congruent and incongruent trials (likelihood ratio test p = 0.214). We did not observe effects of S1 congruency on measures of frontal theta or motor preparation responses (Figures 5C, 5D) and Mu/Beta amplitudes were very similar across conditions at the time of S2 target onset.

**Figure 5.**
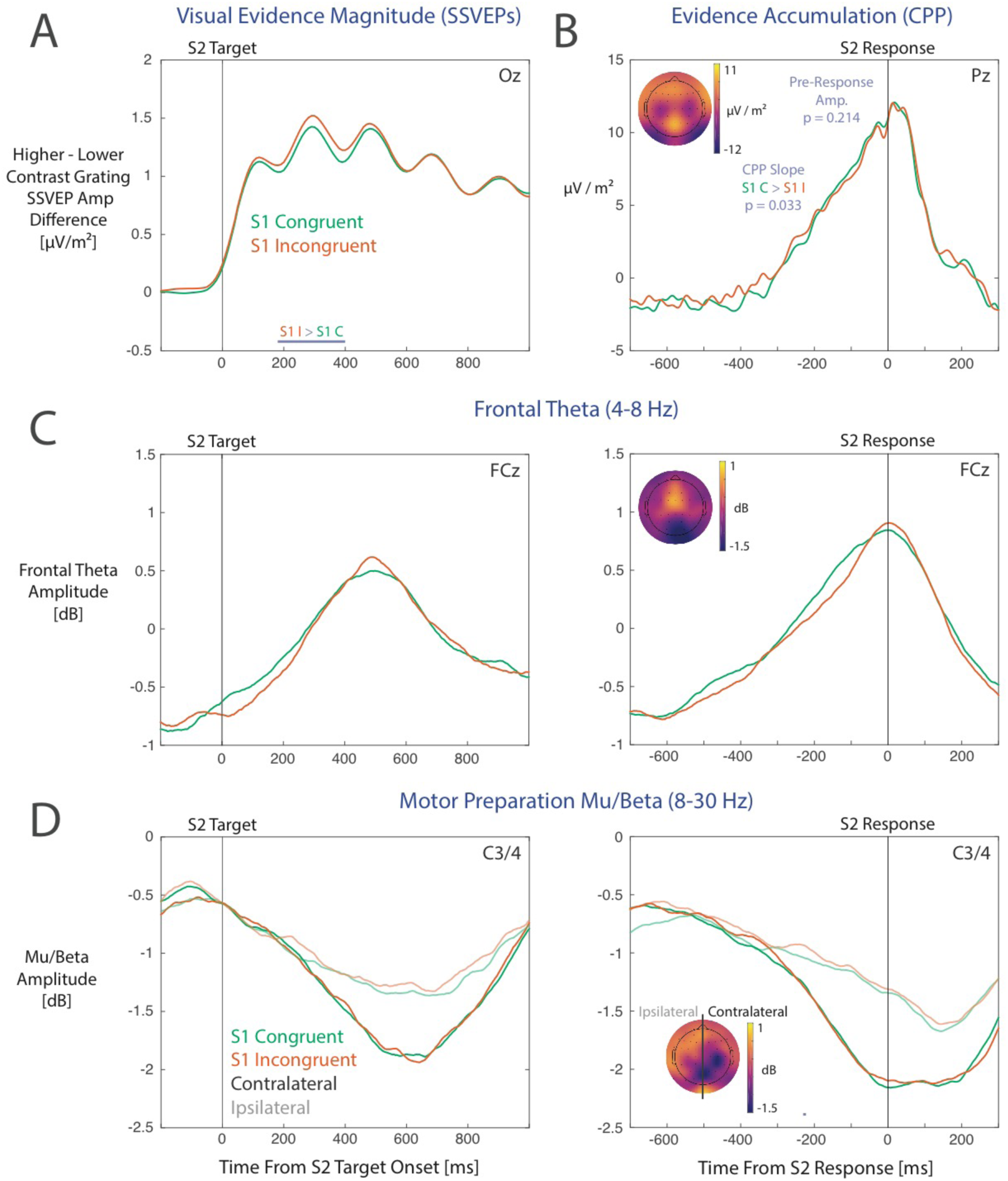
Neural responses following S2 targets by S1 congruency. A) SSVEP difference measures indexing the magnitude of evidence represented in visual cortex in favour of the dominant (i.e. higher contrast) target grating. B) CPP amplitudes leading up to the keypress response. C) Frontal theta amplitudes relative to the target onset time (left panel) and time of response (right panel). D) Mu/Beta amplitudes indexing motor preparation for electrodes contralateral and ipsilateral to the response hand. Scalp maps depict the average magnitudes of each signal over the 200 ms period preceding the keypress response, averaged across S1 congruent and incongruent conditions.

#### 3.3.2 Effects of S2 evidence strength

Effects of S2 evidence strength were clearly visible across all neural measures. Sensory evidence magnitudes scaled with the contrast differences between higher and lower contrast gratings (i.e. evidence strength) from ∼0-800 ms from S2 target onset (Figure 6A). The build-up rate of the CPP was also faster in trials with higher evidence strength, F(2, 52) = 12.02, p < 0.001 (Figure 6B) and pre-response CPP amplitudes were also smaller in weaker evidence strength conditions (likelihood ratio test p = 0.020). Mu/Beta motor preparation amplitude profiles also followed a similar pattern to the CPP, with steeper (negative-going) build-up rates for stronger evidence strengths leading up to the response at contralateral electrodes, but similar amplitudes around the time of response (Figure 6D).

**Figure 6.**
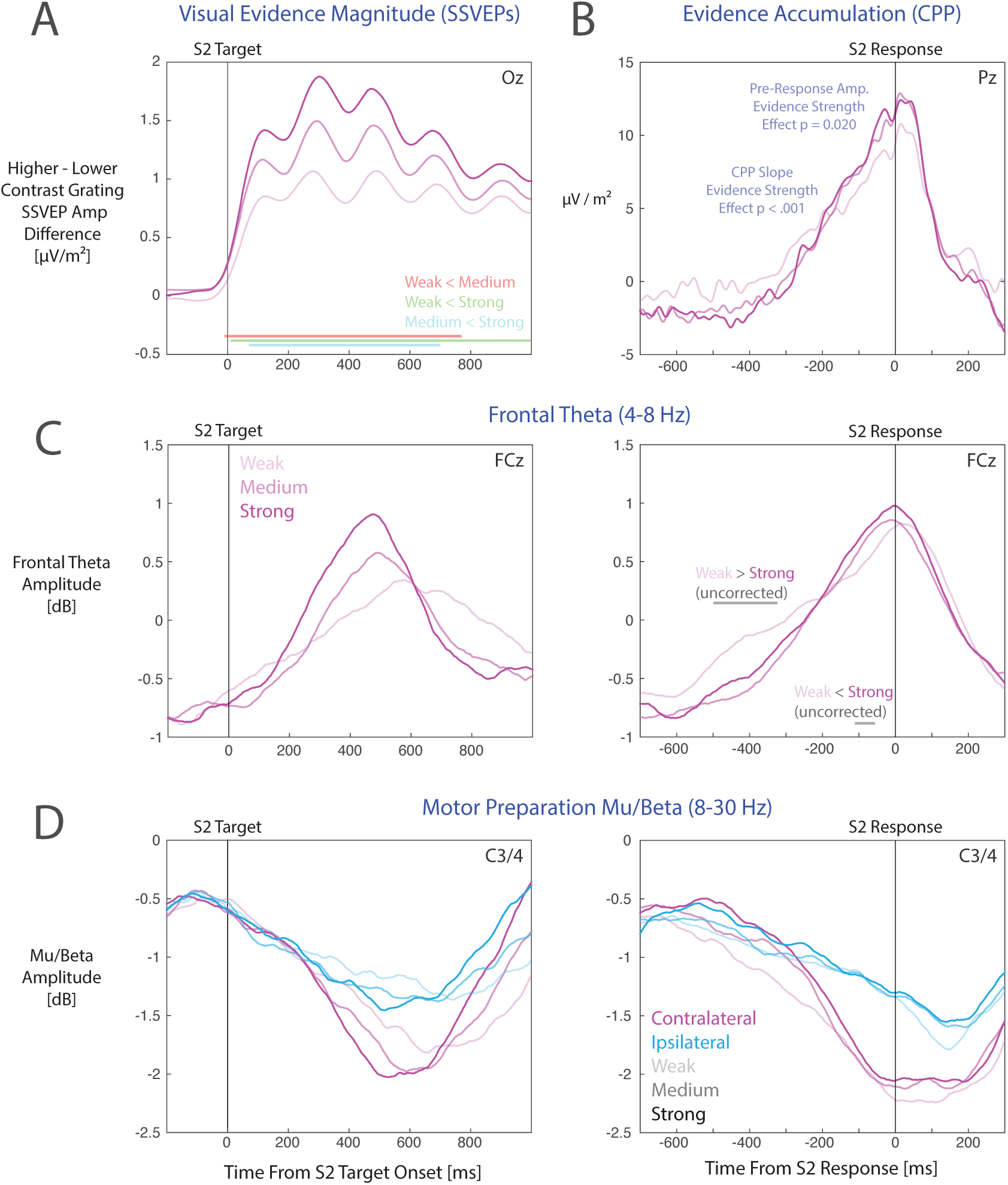
Neural responses following S2 targets by S2 evidence strength. A) SSVEP difference measures indexing the magnitude of evidence represented in visual cortex in favour of the dominant (i.e. higher contrast) target grating. Coloured bars denote time points at which there were statistically significant differences across conditions. B) CPP amplitudes leading up to the keypress response. C) Frontal theta amplitudes relative to the target onset time (left panel) and time of response (right panel). D) Mu/Beta amplitudes indexing motor preparation for electrodes contralateral and ipsilateral to the response hand.

Similar to motor preparation profiles following S1 targets, there appeared to be a gradual build-up of Mu/Beta amplitudes at both contralateral and ipsilateral electrodes from 0-300 ms following S2 target onset. After this point, motor preparation trajectories were determined by the evidence strength of the S2 target. However, motor preparation amplitudes preceding S2 target onset were much less negative-going than those preceding the S1 target (approx. −0.5 dB compared to −1.2 dB, relative to a −2 dB motor execution threshold), indicating relatively less preparatory motor activity preceding the onset of the S2 target.

#### 3.3.3 Frontal theta amplitude profiles

Frontal theta amplitude profiles for S2 targets differed markedly from those profiles observed following S1 targets. Theta amplitudes for S2 targets did not exhibit the rise-until-response pattern that was seen for S1 targets. Instead, theta amplitudes following S2 targets more closely resembled the trajectories of CPP and Mu/Beta amplitudes (Figure 6C left panel; for time-frequency plots see Supplementary Figure S4). Notably, frontal theta amplitudes in the time window immediately preceding the response did not appear to be larger in trials with weaker evidence strength and by association slower RTs (Figure 6C right panel). To investigate this, we performed post-hoc mass-univariate comparisons of response-locked theta amplitudes across strong and weak evidence strength conditions (corresponding to trials with fast and slow RTs). Larger theta amplitudes were observed in weak evidence strength conditions between −500 to −300 ms preceding the response (uncorrected for multiple comparisons), associated with a less steep theta amplitude build-up rate prior to the keypress response. However, these differences were not statistically significant when applying a cluster-based correction for multiple comparisons. Notably, Frontal theta amplitudes were not significantly larger in weak evidence conditions from −300 ms to 50 ms relative to response onset, which is where the bulk of the effects in S1 response-locked epochs were found. Instead, amplitudes in the weak evidence trials tended to be smaller than in strong evidence trials.

In sum, the qualitative patterns of covariation between frontal theta, CPP and Mu/Beta amplitudes differed across S1 and S2 task phases. Following S1 targets, there were larger pre-response theta amplitudes in trials with slower RTs, due to a pattern whereby theta amplitudes gradually rose at a steady rate until the time of the response (see also Tollner et al., 2017). This also produced larger theta amplitudes for incongruent S1 stimuli, which co-occurred with smaller CPP amplitudes and highly similar Mu/Beta amplitudes prior to the S1 response. In contrast, there were no clear theta-RT correlations for S2 targets, and theta-band activity closely resembled the amplitude profiles of the CPP and Mu/Beta activity (see also van Vugt et al., 2012).

## 4. Interim Discussion

We found evidence for two adjustments that occur following exposure to incongruent stimuli: A temporary boost in the magnitude of sensory evidence represented in visual cortex, indexed by effects on SSVEPs, and a slowing of evidence accumulation rates, indexed by shallower CPP slopes. Here we note that, although CPP slopes significantly differed by S1 congruency, the ERP waveforms in the group-averaged plots (in Figure 5B) were highly similar, and we did not observe statistically significant differences in RTs by S1 congruency.

One likely reason for why the observed effects on both the CPP and RTs were small is that we only presented congruent stimuli at S2. Congruency sequence effects are sometimes (but not always) smaller for congruent compared to incongruent Flanker stimuli (Mayr et al., 2003; Nieuwenhuis et al., 2006; but see Gratton et al., 1992; Duthoo et al., 2013). Another possibility is that distinct adjustments occurred at different stages of the decision process (e.g., as found by Steinemann et al., 2018), including a transient RT speeding effect seen over similar response-stimulus intervals (Egner et al., 2010) and a slowing of evidence accumulation, which would have cancelled-out any large RT effects that might be seen for S2 congruent stimuli. Accordingly, we ran a behavioural follow-up experiment that also presented incongruent stimuli at S2, which allowed us to further test the hypotheses that i.) evidence accumulation rates are reduced following S1 incongruent stimuli, and ii.) the reduction in evidence accumulation rates is associated specifically with the sensory information provided by the distractor element of the modified flanker stimuli.

The latter hypothesis is associated with theorised shifts of attention away from the distractor in conflict monitoring models (e.g., Shenhav et al., 2013). Assuming that evidence accumulation rates reflect the sum of sensory evidence provided by the target and distractor stimuli (e.g., as postulated in contemporary evidence accumulation models designed for conflict task data, such as the Shrinking Spotlight Model; White et al., 2011, 2018), this would predict that accumulation rates would be relatively faster for S2 incongruent stimuli when S1 was also incongruent (due to reduced evidence weighting of the unhelpful distractor), but slower for S2 congruent stimuli (whereby the distractor would provide evidence for the correct response). Importantly, changes in the rate of evidence accumulation would also predict larger congruency sequence effects on RTs in trials with slower overall responses, indexed by shallower slopes of the delta functions for trials in which S1 was incongruent.

## 5. Experiment 2

To further test for changes in evidence accumulation rates following response conflict, we conducted a subsequent behavioural experiment (N = 29, 21 female, 8 male, aged between 18-37, with 1 participant excluded from analyses due to poor task performance). The trial structure was identical to the original experiment, except that incongruent stimuli were also presented at S2, and that all S2 targets were of medium evidence strength (for full methods and results see the Supplementary Material).

We observed the hypothesised pattern of effects. Differences in behavioural responses to S1 targets by S1 congruency were consistent with the first experiment (Figures 7A-C). For RTs following S2 targets, there was an S1 congruency by S2 congruency interaction effect (likelihood ratio test p < 0.001) reflecting the typical pattern of congruency sequence effects (e.g., Gratton et al., 1992; Duthoo et al., 2013). Participants were faster when responding to S2 incongruent stimuli when S1 stimuli were also incongruent (likelihood ratio test p’s < 0.001). For RTs to S2 congruent stimuli, there was little evidence of an effect of S1 congruency (likelihood ratio test p = 0.620). Participants responded with similar accuracy across S1 congruency conditions (see Figures 7D-E).

**Figure 7.**
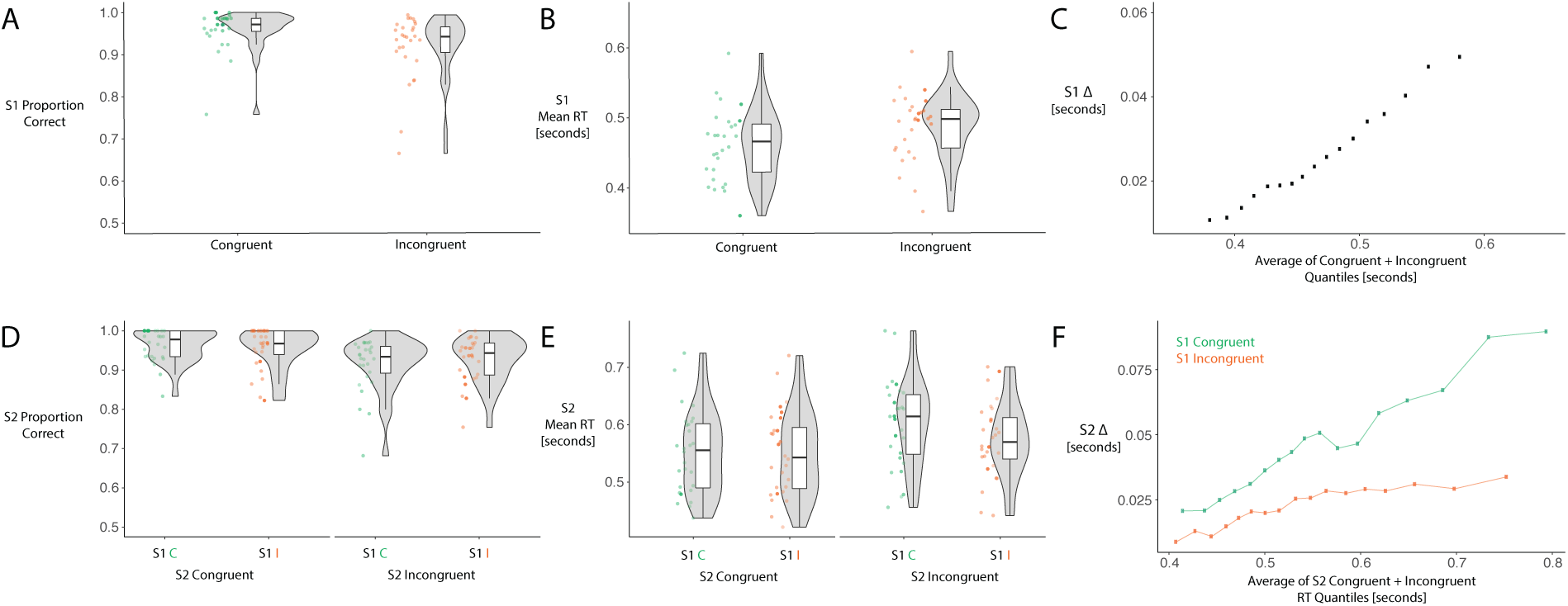
Accuracy and mean RTs following S1 and S2 targets for the behavioural follow-up experiment. For full methods and results see the Supplementary Material. A) Proportions of correct responses following S1 targets with congruent and incongruent distractors. B) Mean RTs for correct responses to S1 targets with congruent and incongruent distractors. C) Delta plot displaying the [incongruent – congruent] differences in RTs for 10-90% quantiles in steps of 5% on the Y axis and the average of each congruent and incongruent condition RT quantile on the X axis. D) Proportions of correct responses to S2 targets split by S1 and S2 congruency. E) Mean RTs for correct responses to S2 targets, split by S1 and S2 congruency. F) Delta plots displaying the S2 [incongruent – congruent] differences for each RT quantile, plotted separately for S2 targets following S1 congruent and S1 incongruent stimuli. Note that the slope of the delta function is shallower for trials whereby S1 was incongruent.

The S2 stimulus delta plots for S1 congruent and incongruent conditions revealed a clearly visible reduction in the slope of the delta function when S1 was incongruent (Figure 7F). For example, when the S1 target was congruent, the incongruent distractors in the S2 stimuli slowed RTs to a much larger degree in trials with overall slower (e.g., > 0.6 second) RTs, compared to trials with faster RTs. This pattern is typical of Flanker tasks (Ulrich et al., 2015). However, this effect was much less pronounced (i.e. the delta function was substantially shallower) when S1 was incongruent. Under the assumption that evidence accumulation rates reflect a weighed sum of the sensory evidence provided by the target and distractor elements (e.g., in White et al., 2011, 2018), this pattern of RTs is consistent with a down-weighting of the influence of the distractor in determining the evidence accumulation rate within a trial.

## 6. General Discussion

To characterise the adjustments to decision-making processes that are implemented following response conflict, we recorded EEG during a modified Flanker task and traced the temporal dynamics of sensory evidence representation (measured using SSVEPs), decision evidence accumulation (inferred from slope and peak measures of the CPP component), preparatory motor activity (measured as Mu/Beta activity amplitudes) and frontal theta (O’Connell et al., 2012; Cohen & Donner, 2013; Murphy et al., 2015; Steinemann et al., 2018). We identified neural markers of two distinct adaptations following exposure to incongruent stimuli, including a boost in the magnitude of sensory evidence represented in visual cortex and a small but statistically significant slowing of the rate of decision evidence accumulation. Evidence consistent with reductions in evidence accumulation rates following incongruent stimuli was also found in a foll0w-up behavioural experiment, which also showed that this effect related to sensory evidence provided by the incongruent (i.e. distracting) stimulus elements. Taken together, our findings provide initial evidence that shifts of attention in conflict monitoring accounts are implemented as adjustments to evidence accumulation rates, rather than other effects which might be conceptually associated with attention.

In addition, we identified ramping motor activity at electrodes both contralateral and ipsilateral to the response hand that started before S1 target onset (Figure 4D left panel) and was visible following S2 onset. These patterns of motor activity have been previously linked to decision urgency signals (Steinemann et al., 2018; Kelly et al., 2020) that can influence the timing of behavioural responses under similar conditions to most existing conflict tasks (e.g., Cohen & Donner, 2013; Tollner et al., 2017). We also observed markedly different frontal theta (4-8 Hz) activity profiles across S1 and S2 task phases, corresponding to temporal profiles reported in response conflict tasks (e.g., Cohen & Donner, 2013; Tollner et al., 2017) and those observed in other perceptual decision tasks (e.g., van Vugt et al., 2012). Based on these frontal theta amplitude profiles, and their qualitative patterns of covariation with neural markers of evidence accumulation and motor activity, we discuss some potential contributions to frontal theta amplitude measures that may be distinct from effects that are specifically associated with response conflict.

### 6.1 Adjustments to Decision Processes Following Response Conflict

We found typical effects of stimulus congruency on behavioural and neural responses to S1 targets in our modified Flanker task, consistent with existing work (e.g., Gratton et al., 1992; Ulrich et al., 2015; Tollner et al., 2017). More importantly, we also identified neural markers of two distinct adjustments to decision-making processes following response conflict. Firstly, we observed increases in the magnitude of decision-relevant sensory evidence as represented in visual cortex, from approximately 200 ms after S2 target onset. Although this observed increase might seem surprising, the timing of this effect is consistent with effects of increased pupil dilation following response conflict (Geva et al., 2013; van der Wel & van Steenbergen, 2018), which is associated with larger sensory evidence magnitudes in favour of the correct response (as indexed by SSVEPs) in a highly similar task to that in the current study (Steinemann et al., 2018). Similar changes in pupil dilation are also observed following surprising events and decision errors, which have been linked to the orienting response (reviewed in Wessel, 2017). Given that there are similar midfrontal effects on neural activity following conflict, error detection and surprise, which have been incorporated into conflict monitoring-based accounts (Cavanagh & Frank, 2014), the time-course of the orienting response and associated neurophysiological phenomena may be important to consider when modelling congruency sequence effects.

Somewhat counterintuitively, these effects on SSVEP measures co-occurred with slower rates of evidence accumulation, as indexed by changes in the slope of the CPP. Notably, we could detect this effect even though our SSVEP measures of sensory evidence were larger, which is associated with steeper rather than shallower CPP build-up rates (O’Connell et al., 2012; Steinemann et al., 2018). This may reflect a shift of attention away from the inner grating annulus that comprised the S1 distractor stimulus, which is conceptually similar to shifts of attention away from distractor stimuli as described in conflict monitoring models (Botvinick et al., 2004; Shenhav et al., 2013). Our results suggest that this so-called attention shift might be implemented as a change in the accumulation rate of decision evidence, and that this process is not necessarily a downstream effect of changes in the magnitude of sensory evidence representations in visual cortex. However, we caution that the observed size of these effects on the CPP were rather small, and did not co-occur with expected differences in RTs across conditions. Before drawing any strong inferences, our ERP results should be replicated in situations that produce large congruency sequence effects on RTs, for example using the design in Jiang et al. (2018).

Despite these issues, we did find additional evidence consistent with post-conflict changes in evidence accumulation rates in a subsequent behavioural experiment, whereby we also presented incongruent S2 stimuli. Specifying changes in evidence accumulation rates based on the ERP results allowed us to derive predictions regarding how congruency sequence effects will be reflected in patterns of fast, medium and slow RTs. This is in contrast to existing models, which could only generate predictions relating to mean RTs (Botvinick et al., 2001; Shenhav et al., 2013). Based on the notion that post-conflict adjustments specifically reflect a down-weighting of sensory evidence provided by the distractor stimulus elements in the evidence accumulation process, we expected congruency sequence effects on RTs for S2 targets to be progressively larger in trials with slower overall responses. This would be indexed by less steep delta functions (indexing effects of S2 congruency across fast, medium and slow RT quantiles) for trials where the S1 stimulus was incongruent, compared to when S1 was congruent. We observed precisely this pattern of effects, lending further support to the notion that post-conflict adjustments are associated with changes in how decision evidence is accumulated over time. This indicates that cognitive control processes described in conflict monitoring theories (e.g., Botvinick et al., 2001; Shenhav et al., 2013) exert influence over how sensory evidence is weighted at the evidence accumulation stage of decision-making. It also suggests that the concept of attention shifts in these models does not necessarily correspond to shifts in visual attention as understood through, for example, normalisation models of attention (e.g., Reynolds & Heeger, 2009). We propose that the concept of ‘attention’ in these accounts should be revised to explicitly describe the decision-processes that are influenced following conflict.

Interestingly, our findings of changes in evidence accumulation rates broadly agree with mathematically-formalised evidence accumulation models that were developed specifically to describe decision processes in conflict tasks (Hübner et al., 2010; White et al., 2011, 2018; Ulrich et al., 2015). Each of these models describes effects of conflict as influencing the (nonstationary) trajectory of the decision variable, leading to changes in trial-averaged drift rates, rather than effects on other parameters such as sensory encoding duration or decision thresholds. Our findings suggest that such models can be extended to capture congruency sequence effects by changing the weightings of the target and distractor elements in determining the path of the decision variable following response conflict.

One notable limitation related to our SSVEP measures is that they captured the pooled visual responses to each grating orientation across target and distractor elements of the stimulus. This means that we could not identify reductions in visual responses to the distractor (i.e. shifts of spatial attention) that may have co-occurred with pupil dilation-related increases in SSVEP amplitudes. Future experiments could frequency tag responses to target and distractor elements separately, in order to better dissociate the dynamic allocation of spatial attention from effects related to pupil dilation. Alternatively, future experiments could frequency tag responses to left- and right-tilted gratings for target and distractor elements separately, however this may be difficult to achieve without substantial harmonic overlap within frequency bins when performing discrete Fourier transforms.

Another caveat is that we did not observe a slowing of responses to S2 stimuli following incongruent S1 stimuli in the EEG experiment, which differs from reports of post-conflict slowing effects in many (but not all) previous studies (e.g., Gratton et al., 1992; Mayr et al., 2003; Duthoo et al., 2013; Egner et al., 2010). One possibility is that participants learned to expect only congruent stimuli at S2, which reduced the prevalence of strategic adjustments after exposure to response conflict at S1. However, we also did not observe post-conflict slowing effects for congruent S2 stimuli in the follow-up experiment whereby participants could expect both congruent and incongruent S2 stimuli. Another possibility is slowed evidence accumulation co-occurred with other arousal-related effects that led to a general speeding of RTs (as reported by Egner et al., 2010), where the RT measures in our models reflected the sum of these counter-acting effects. Based on these findings we advise caution when directly comparing behaviour in our modified Flanker task to those from more typical Flanker tasks. We also caution that our EEG effects pertain specifically to the consequences of exposure to an incongruent stimulus, and do not capture co-occurring effects of expectancies of an upcoming incongruent stimulus that may be present in existing studies. Given that others have reported distinct and additive effects of prior exposure and expectancy (Alpay et al., 2009), future work could additionally characterise the behavioural and ERP correlates of each of these factors.

The EEG analysis framework used here could also be adapted to further develop models of other phenomena related to cognitive control, such as proportion congruent effects (reviewed in Schmidt, 2019) and slowing of responses in trials following low-confidence decisions (Desender et al., 2019).

### 6.2 Effects of Ramping Motor Activity

We also observed patterns of Mu/Beta activity that may be informative for modelling decision-making performance in conflict tasks, particularly in situations when strict response deadlines are enforced. As recently reported by Steinemann et al. (2018) and Kelly et al. (2019), we also found gradually building motor activity during both S1 and S2 task phases, indexed by negative-going Mu/Beta spectral amplitudes at electrodes both contralateral and ipsilateral to the response hand. This motor activity started building even before the S1 target was presented, and a behavioural response (i.e. a keypress) was made when it reached a fixed threshold (see also Donner et al., 2009; O’Connell et al., 2012). Notably, the extent of decision evidence accumulation (indexed by the CPP) reached lower levels in trials with slower RTs, indicating that this build-up of evidence-independent motor activity forced keypress responses based on less decision evidence in these trials, as compared to trials with faster RTs. This gradual build-up of decision evidence-independent motor activity can be understood as the neural implementation of decision thresholds that collapse toward zero over the course of each task phase (Ditterich, 2006; Cisek et al., 2009; Drugowitsch et al., 2012). The notion of collapsing thresholds is highly controversial in the behavioural modelling literature (Ratcliff et al., 2016). Modelling results have favoured collapsing thresholds only when there are strict response deadlines or strong pressure to respond quickly (Hawkins et al., 2015; Voskuilen et al., 2016; Evans et al., 2019). However, neurophysiological investigations in primates have consistently found support for collapsing thresholds implemented within motor areas (e.g., Ditterich, 2006; Cisek et al., 2009; Heitz & Schall, 2012; Murphy et al., 2016; Steinemann et al., 2018).

In our experiment there was a critical difference between S1 and S2 task phases that determined how much this ramping motor activity influenced the timing of behavioural responses. During the S1 task phase, when there was a strict (800 ms) response deadline, the (negative-going) Mu/Beta amplitudes indexing motor activity had traversed almost halfway to the motor execution threshold by the time of S1 target onset, meaning that even small additional effects of ramping motor activity could trigger a keypress response following S1 targets. However, this was not the case during the S2 task phase, whereby the level of motor activity was further from the threshold at S2 target onset (see also Steinemann et al., 2018; Kelly et al., 2020). Consequently, Mu/Beta amplitude trajectories were largely determined by trajectories of decision evidence accumulation from around 300 ms following S2 target onset.

Strict response deadlines are often used in response conflict tasks (e.g. Cohen & Donner, 2013; Cohen & Cavanaugh, 2011), and this may be why RT distributions in these tasks have been difficult to account for using standard diffusion models with time-invariant decision thresholds (see Ulrich et al., 2015; Servant et al., 2016). Modelling effects of ramping motor activity (as collapsing decision bounds) may improve model fits in these situations (e.g., Kelly et al., 2020), and should be considered when developing computational models of response conflict effects (for a review of contemporary models see White et al., 2018).

We also note that the extent of Mu/Beta at contralateral and ipsilateral electrodes to the response hand did not appear to significantly differ by the presence/absence of response conflict in the S1 stimulus. This may appear as evidence against conflict monitoring accounts that presuppose differences in ipsilateral motor activity for incongruent stimuli. However, it is unclear whether Mu/Beta activity is a direct correlate of those motor-related signals that provide input to medial prefrontal cortex in conflict monitoring accounts (e.g., Cohen, 2014). To better elucidate the role of Mu/Beta-related motor action preparation in relation to conflict monitoring it may be informative to trace the time-course of these oscillatory signals in other conflict tasks (e.g., Cohen & van Gaal, 2014; Tollner et al., 2017; Jiang et al., 2018).

### 6.3 Observed Contributions to Frontal Theta Amplitude Measures

By tracing the temporal profiles of frontal theta amplitudes across S1 and S2 task phases, we also provide preliminary evidence for two contributions to frontal theta amplitude measures that occur in addition to effects of response conflict detection. We note here that these are based on visual inspection of the data, and should ideally be replicated and further characterised in future work.

The first source of frontal theta appeared to track motor activity associated with a task-related motor action, as quantified using Mu/Beta spectral amplitudes in our experiment. This resembled the build-up of decision evidence, which is likely why frontal theta has been previously proposed as a neural correlate of evidence accumulation (van Vugt et al., 2012; Werkle-Bergner et al., 2014). However, our results suggest that this source of frontal theta correlates with motor activity rather than decision evidence accumulation (as in the model of Brown & Braver, 2008). This is because both theta and motor activity began to increase before the onset of the S1 target (Figures 4C, 4D) before decision evidence would begin accumulating in our task. This assumption could be further tested by assessing whether accumulation-to-bound frontal theta dynamics are also found in perceptual decision tasks that do not require motor responses (e.g., O’Connell et al., 2012).

The second source of frontal theta relates to the response deadline used in our task, where response deadlines are also present in most existing EEG-based studies of conflict (Cohen & Cavanaugh, 2011; Cohen & Donner, 2013; Cohen & van Gaal, 2014; Jiang et al., 2018). In previous studies involving conflict tasks, researchers have reported a steady increase in theta power at a fixed rate over the course of each trial that stopped rising ∼100 ms prior to a keypress response (Cohen & Cavanaugh, 2011; Cohen & Donner, 2013; Cohen & van Gaal, 2014; Jiang et al., 2018; see Figure 6C in Tollner et al., 2017) and we also observed this following S1 targets. This dynamic produces larger theta amplitudes immediately preceding behavioural responses in trials with longer RTs. However, following S2 targets, and in a previous study that did not use a response deadline (van Vugt et al., 2012) frontal theta amplitudes more closely tracked the build-up of decision evidence, and rose at a faster rate following stimulus onset in trials with faster RTs (van Vugt et al., 2012).

Here we present two potential explanations for this pattern of theta amplitude profiles. The first explanation is that, in our experiment, the rate of evidence accumulation was slower in trials with incongruent S1 stimuli, as evidenced by the shape of the delta plot and differences in CPP build-up rates. At the same time, motor activity gradually increased over the S1 task phase at a similar rate for both S1 congruent and incongruent stimulus conditions, which can be attributed to ramping effects of evidence-independent motor urgency (Steinemann et al., 2018; Kelly et al., 2020). Because the extent of motor activity is determined by both the accumulation of decision evidence and urgency effects (Kelly et al., 2020), the growing influence of urgency over time would lead to levels of motor activity that exceed that which is normally associated with the current level of accumulated decision evidence. During the S1 task phase this discrepancy between the amount of decision evidence and the extent of motor activity would have grown larger as time passes, and would be larger at the time of the keypress response in trials with incongruent stimuli and slower RTs. If frontal theta amplitudes track the magnitude of this discrepancy, it would lead to the same rise-until-response dynamics observed in our results, as well as a positive correlation between pre-response theta amplitudes and RTs. Frontal theta is known to correlate with a range of prediction error signals that index a mismatch between expected and actual behavioural outcomes, rewards, and other decision-relevant features (reviewed in Cavanagh & Frank, 2014). Here, it is also plausible that frontal theta may signal discrepancies between the amount of evidence for a decision and the extent of preparation corresponding to decision-relevant motor actions, which would arguably provide a critical source of information for continuous performance monitoring and error detection (Kelly et al., 2020). These effects could be dissociated from more general ‘time-on-task’ effects (see Tollner et al., 2017) by assessing whether the presence of a strict response deadline is necessary to observe these temporal dynamics. If this reflects a general increase in theta with increasing RT, then the pattern should persist in the absence of a strict response deadline.

The second, and equally likely, source of these theta dynamics is that participants made larger numbers of partial errors when RTs were slower and closer to the response deadline. Partial errors are defined here as when a participant initiates a motor response for the incorrect choice option, captured as small muscle activations using electromyography (EMG), but does not apply enough force to press down the response key, and instead presses the response key for the correct choice instead (Coles et al., 1995; Cohen & van Gaal, 2014). Partial errors are associated with increased theta power in the time window preceding the keypress (e.g., Cohen & van Gaal, 2014) and it is reasonable to expect that higher proportions of trials may contribute to trial-averaged theta measures when RTs are slower. As we did not record EMG data, we could not assess the prevalence of partial errors across conditions in our study, but we recommend that others record EMG when employing conflict tasks in future work.

Here it is important to note that theta-band activity as measured in our experiment likely reflects both EEG signals that are phase-locked and non-phase-locked relative to the stimuli or keypress responses. Researchers have attempted to isolate and subtract phase-locked contributions to theta-band activity by subtracting trial-averaged ERPs prior to time-frequency power estimation (e.g., Cohen & Donner, 2013). However, this approach relies on the assumption that phase-locked EEG signals are of identical amplitude at each time point relative to the event of interest. Trial-by-trial variations in the amplitudes of phase-locked signals would not be perfectly subtracted, and so would presumably contribute to theta-band estimates. Consequently, our findings more directly relate to how theta power is measured in similar designs, rather than the underlying generators of phase-locked and non-phase-locked frontal theta power.

### 6.4 Conclusion

We traced the neural dynamics of multiple processes that are critical for perceptual decision-making, and characterised two distinct adjustments to decision processes that occur following response conflict. We report effects on how stimuli are processed in the visual system, indexed by post-conflict increases in SSVEP amplitudes. We also uncovered evidence of drift rate adjustments, whereby sensory evidence provided by the distractor is down-weighted in determining the trial-averaged drift rate. Our findings help to specify how adjustments to decision-making processes are implemented after encountering conflicting stimulus information, which is critical for extending and better specifying contemporary models of cognitive control.

## Acknowledgements

We thank Redmond O’Connell and Patrick Cooper for their feedback and advice regarding a previous version of this manuscript. This project was supported by an Australian Research Council Grant (ARC DP160103353) to S.B. and R.H. Funding sources had no role in study design, data collection, analysis or interpretation of results.

