## Supplementary Material for "Tracking dynamic adjustments to decision making and performance monitoring processes in conflict tasks"

### Supplementary Material List of Contents

1. Mixed effects regression model structures and coefficients for analyses of accuracy and response times (RTs)
2. Mixed effects regression model structures for analyses of centro-parietal positivity (CPP) pre-response mean amplitudes
3. Numbers of epochs in each condition that were included in EEG analyses
4. Mean RTs for correct and error responses to S1 and S2 targets
5. Time-Frequency Plots for S1 Targets at Electrode FCz
6. Lateralised [Contralateral – Ipsilateral] Mu/Beta Differences for S1 Targets
7. Time-Frequency Plots for S2 Targets at Electrode FCz
8. Supplementary Experiment 2 Methods
9. Supplementary Experiment 2 Results
10. Supplementary material reference list

### 1. Mixed Effects Regression Model Structures and Coefficients for Analyses of Accuracy and RTs

Code for fitting all models listed below will be available at https://osf.io/eucqf/ at the time of publication. To test for fixed effects of each variable of interest on accuracy and RT measures, we compared models with and without the fixed effect of interest, but with identical random effects structures, using likelihood ratio tests. Models were fit using the R package lme4 (Bates et al., 2015). We used a random effects structure that included a random intercept by participant and random slopes of all variables that would conceivably influence accuracy rates and RTs, except where adding these prevented the model from converging. For S1 responses these variables included S1 target orientation (left, right) and S1 congruency (congruent, incongruent). For S2 responses these variables included S1 congruency (congruent, incongruent), S2 evidence strength (weak, medium, strong), and the additive and interactive effects of S1 and S2 orientation (left, right) to account for differences across 15 Hz and 20 Hz target grating contrast reversal rates and any effects of S1/S2 target orientation repetition or alternation. To allow the models to converge, when testing for effects of S1 congruency on S2 accuracy and RTs, we retained only the random slope of S1 congruency, and excluded random slopes of S2 evidence strength and S1 and S2 orientation.

### 1.1 List of Variable Names and Definitions

`RT_S1` – The response time for S1 targets in seconds

`RT_S2` – The response time for S2 targets in seconds

`outcome_S1` – The decision outcome following S1 targets (correct or error)

`outcome_S2` – The decision outcome following S2 targets (correct or error)

`congruency` – Whether the S1 target and distractor was congruent or incongruent

`orientation_S1` – The direction of higher contrast gratings in the S1 target stimulus (left or right)

`orientation_S2` – The direction of higher contrast gratings in the S2 target stimulus (left or right)

`evidence_strength` – The contrast difference between higher and lower contrast gratings for S2 targets (weak, medium or strong)

`ID` – The participant ID number (used for defining random intercepts and slopes)

### 1.2 Regression Model Equations and Coefficients

Full model equations are listed below. Here, we follow the conventions of the lme4 package notation (Bates et al., 2015) when describing the structure of mixed effects regression models. Model coefficients for the full model (including the fixed effect of interest) are displayed below each set of model equations.

#### 1.2.1 Testing for an Effect of S1 Congruency on S1 RTs

For analyses of S1 RTs, we fit generalised linear mixed effects regression models (Gamma family) with an identity link function. We compared the following model including effects of S1 congruency:

```
RT_S1 ~ orientation_S1 + congruency + (1 + orientation_S1 +
congruency | ID)
```

With a comparison model that did not include a fixed effect of congruency:

```
RT_S1 ~ orientation_S1 + (1 + orientation_S1 + congruency | ID)
```

Coefficients for the generalised linear mixed model (Gamma family, identity link function) fit by maximum likelihood

| | | | | | |
|---|---|---|---|---|---|
| | AIC | BIC | Log Likelihood | Deviance | |
| | -24050.9 | -23977.0 | 12035.5 | -24070.9 | |
| Scaled Residuals: | | | | | |
| | Min | 1Q | Median | 3Q | Max |
| | -4.308 | -0.638 | -0.089 | 0.556 | 5.460 |
| Random Effects: | | | | | |
| | Groups | Name | Variance | Std. Dev | |
| | ID | (Intercept) | 0.0004 | 0.021 | |
| | | S1 Orientation | 0.0002 | 0.016 | |
| | | S1 Congruency | 0.0004 | 0.022 | |
| | Residual | | 0.033 | 0.183 | |
| Number of observations: 11960, groups: ID, 29 | | | | | |
| Fixed Effects | | | | | |
| | | Estimate | Std. Error | t value | |
| | (Intercept) | 0.461 | 0.012 | 37.499 | |
| | S1 Orientation | 0.013 | 0.007 | 1.911 | |
| | S1 Congruency | 0.066 | 0.011 | 6.221 | |

#### 1.2.2 Testing for an Effect of S1 Congruency on Proportion Correct Responses to S1 Targets

For analyses of proportion correct responses (i.e., accuracy) for S1 targets, we fit generalised linear mixed effects regression models (Binomial family) with a logit link function. We compared the following model including effects of S1 congruency:

```
outcome_S1 ~ orientation_S1 + congruency + (1 + orientation_S1 +
congruency | ID)
```

With a comparison model that did not include a fixed effect of congruency:

```
outcome_S1 ~ orientation_S1 + (1 + orientation_S1 + congruency | ID)
```

Coefficients for the generalised linear mixed model (Binomial family, logit link function) fit by maximum likelihood

| | | | | | |
|---|---|---|---|---|---|
| | AIC | BIC | Log Likelihood | Deviance | |
| | 6285.8 | 6353.0 | 6267.8 | 6267.8 | |
| Scaled Residuals: | | | | | |
| | Min | 1Q | Median | 3Q | Max |
| | -11.091 | 0.138 | 0.193 | 0.318 | 0.727 |
| Random Effects: | | | | | |
| | Groups | Name | Variance | Std. Dev | |
| | ID | (Intercept) | 0.790 | 0.889 | |
| | | S1 Orientation | 0.383 | 0.618 | |
| | | S1 Congruency | 0.532 | 0.729 | |
| Number of observations: 12973, groups: ID, 29 | | | | | |
| Fixed Effects | | | | | |
| | | Estimate | Std. Error | z value | |
| | (Intercept) | 3.895 | 0.195 | 20.023 | |
| | S1 Orientation | -0.254 | 0.141 | -1.799 | |
| | S1 Congruency | -1.595 | 0.170 | -9.362 | |

#### 1.2.3 Testing for an Effect of S2 Evidence Strength on Proportion Correct Responses to S2 Targets

For analyses of proportion correct responses (i.e., accuracy) for S2 targets, we fit generalised linear mixed effects regression models (Binomial family) with a logit link function. We compared the following model including effects of S2 evidence strength:

```
outcome_S2 ~ orientation_S1 * orientation_S2 + evidence_strength +
(1 + evidence_strength | ID)
```

With a comparison model that did not include a fixed effect of S2 evidence strength:

```
outcome_S2 ~ orientation_S1 * orientation_S2 +
(1 + evidence_strength | ID)
```

Coefficients for the generalised linear mixed model (Binomial family, logit link function) fit by maximum likelihood

| | | | | | |
|---|---|---|---|---|---|
| | AIC | BIC | Log Likelihood | Deviance | |
| | 5125.3 | 5213.9 | -2550.7 | 5101.3 | |
| Scaled Residuals: | | | | | |
| | Min | 1Q | Median | 3Q | Max |
| | -15.854 | 0.116 | 0.185 | 0.268 | 1.120 |
| Random Effects: | | | | | |
| | Groups | Name | Variance | Std. Dev | |
| | ID | (Intercept) | 0.497 | 0.705 | |
| | | S2 Evidence Strength (Linear) | 0.304 | 0.552 | |
| | | S2 Evidence Strength (Quadratic) | 0.066 | 0.257 | |
| Number of observations: 11890, groups: ID, 29 | | | | | |
| Fixed Effects | | | | | |
| | | Estimate | Std. Error | z value | |
| | (Intercept) | 2.445 | 0.148 | 16.497 | |
| | S1 Orientation | 1.536 | 0.129 | 11.931 | |
| | S2 Orientation | 1.358 | 0.125 | 10.902 | |
| | S2 Evidence Strength (Linear) | 0.859 | 0.130 | 6.584 | |
| | S2 Evidence Strength (Quadratic) | -0.303 | 0.098 | -3.093 | |
| | S1 Orientation * S2 Orientation | -2.972 | 0.179 | -16.633 | |

##### 1.2.4 Testing for an Effect of S1 Congruency on Proportion Correct Responses to S2 Targets

For analyses of proportion correct responses (i.e., accuracy) for S2 targets, we fit generalised linear mixed effects regression models (Binomial family) with a logit link function. We compared the following model including effects of S1 congruency:

```
outcome_S2 ~ congruency + evidence_strength + orientation_S1 *
orientation_S2 + (1 + congruency | ID)
```

With a comparison model that did not include a fixed effect of S1 congruency:

```
outcome_S2 ~ evidence_strength + orientation_S1 * orientation_S2 +
(1 + congruency | ID)
```

Coefficients for the generalised linear mixed model (Binomial family, logit link function) fit by maximum likelihood

| | | | | | |
|---|---|---|---|---|---|
| | AIC | BIC | Log Likelihood | Deviance | |
| | 5161.3 | 5235.1 | -2570.6 | 5141.3 | |
| Scaled Residuals: | | | | | |
| | Min | 1Q | Median | 3Q | Max |
| | -13.760 | 0.120 | 0.186 | 0.288 | 1.248 |
| Random Effects: | | | | | |
| | Groups | Name | Variance | Std. Dev | |
| | ID | (Intercept) | 0.554 | 0.744 | |
| | | S1 Congruency | 0.005 | 0.073 | |
| Number of observations: 11890, groups: ID, 29 | | | | | |
| Fixed Effects | | | | | |
| | | Estimate | Std. Error | z value | |
| | (Intercept) | 2.458 | 0.159 | 15.427 | |
| | S1 Congruency | -0.062 | 0.087 | -0.711 | |
| | S1 Orientation | 1.523 | 0.129 | 11.841 | |
| | S2 Orientation | 1.347 | 0.124 | 10.828 | |
| | S2 Evidence Strength (Linear) | 0.958 | 0.071 | 13.500 | |
| | S2 Evidence Strength (Quadratic) | -0.215 | 0.072 | -2.995 | |
| | S1 Orientation * S2 Orientation | -2.946 | 0.178 | -16.518 | |

#### 1.2.5 Testing for an Effect of S2 Evidence Strength on S2 RTs

For analyses of S2 RTs, we fit generalised linear mixed effects regression models (Gamma family) with an identity link function. We compared the following model including effects of S2 evidence strength:

```
RT_S2 ~ orientation_S1 * orientation_S2 + evidence_strength +
(1 + evidence_strength | ID)
```

With a comparison model that did not include a fixed effect of S2 evidence strength:

```
RT_S2 ~ orientation_S1 * orientation_S2 +
(1 + evidence_strength | ID)
```

Coefficients for the generalised linear mixed model (Gamma family, identity link function) fit by maximum likelihood

| | | | | | |
|---|---|---|---|---|---|
| | AIC | BIC | Log Likelihood | Deviance | |
| | -9807.7 | -9712.7 | 4916.8 | -9833.7 | |
| Scaled Residuals: | | | | | |
| | Min | 1Q | Median | 3Q | Max |
| | -2.908 | -0.600 | -0.157 | 0.408 | 7.615 |
| Random Effects: | | | | | |
| | Groups | Name | Variance | Std. Dev | |
| | ID | (Intercept) | 0.0015 | 0.039 | |
| | | S2 Evidence Strength (Linear) | 0.0013 | 0.036 | |
| | | S1 Evidence Strength (Quadratic) | 0.0002 | 0.014 | |
| | Residual | | 0.0846 | 0.291 | |
| Number of observations: 11036, groups: ID, 29 | | | | | |
| Fixed Effects | | | | | |
| | | Estimate | Std. Error | t value | |
| | (Intercept) | 0.651 | 0.020 | 32.471 | |
| | S1 Orientation | -0.096 | 0.004 | -23.277 | |
| | S2 Orientation | -0.084 | 0.004 | -19.979 | |
| | S2 Evidence Strength (Linear) | -0.161 | 0.015 | -10.705 | |
| | S2 Evidence Strength (Quadratic) | 0.037 | 0.005 | 6.781 | |
| | S1 Orientation * S2 Orientation | 0.201 | 0.006 | 33.811 | |

#### 1.2.6 Testing for an Effect of S1 Congruency on S2 RTs

For analyses of S2 RTs, we fit generalised linear mixed effects regression models (Gamma family) with an identity link function. We compared the following model including effects of S1 congruency:

```
RT_S2 ~ congruency + evidence_strength + orientation_S1 *
orientation_S2 + (1 + congruency | ID)
```

With a comparison model that did not include a fixed effect of S2 evidence strength:

```
RT_S2 ~ evidence_strength + orientation_S1 * orientation_S2 +
(1 + congruency | ID)
```

Coefficients for the generalised linear mixed model (Gamma family, identity link function) fit by maximum likelihood

| | | | | | |
|---|---|---|---|---|---|
| | AIC | BIC | Log Likelihood | Deviance | |
| | -9554.3 | -9473.9 | 4788.1 | -9576.3 | |
| Scaled Residuals: | | | | | |
| | Min | 1Q | Median | 3Q | Max |
| | -2.863 | -0.610 | -0.168 | 0.403 | 7.574 |
| Random Effects: | | | | | |
| | Groups | Name | Variance | Std. Dev | |
| | ID | (Intercept) | 1.352e-03 | 0.037 | |
| | | S1 Congruency | 4.652e-05 | 0.007 | |
| | Residual | | 8.720e-02 | 0.295 | |
| Number of observations: 11036, groups: ID, 29 | | | | | |
| Fixed Effects | | | | | |
| | | Estimate | Std. Error | t value | |
| | (Intercept) | 0.649 | 0.019 | 34.528 | |
| | S1 Congruency | -0.006 | 0.004 | -1.672 | |
| | S1 Orientation | -0.095 | 0.004 | -22.782 | |
| | S2 Orientation | -0.083 | 0.004 | -19.616 | |
| | S2 Evidence Strength (Linear) | -0.150 | 0.003 | -53.255 | |
| | S2 Evidence Strength (Quadratic) | 0.035 | 0.003 | 13.533 | |
| | S1 Orientation * S2 Orientation | 0.200 | 0.006 | 33.162 | |

### 2. Mixed Effects Regression Model Structures for Analyses of Centro-Parietal Positivity (CPP) Pre-response Mean Amplitudes

For analyses of effects of congruency on CPP amplitudes following S1 targets, a model with a fixed effect of congruency, a random slope of congruency by participant, and a random intercept by participant was compared against a model with the same random effects structure (but without the fixed effect of congruency) using a likelihood ratio test. For analyses of effects on pre-response CPP amplitudes following S2 targets, fixed effects included S1 congruency, S2 evidence strength, as well as additive and interactive effects of S1 and S2 target orientation. Models including the fixed effect of interest and all other candidate fixed effects were compared against models without the fixed effect of interest. All models for S2 responses included a random intercept of participant, but no random slopes, due to convergence problems when random slopes were added. Additionally, we tested for the relationship between pre-response CPP amplitudes and RT following S1 stimuli, by comparing models including all fixed effects described above with a model additionally including an effect of RT, and the same random effects structure as specified above. CPP amplitudes and RT measures were scaled and centred prior to fitting all models.

#### 2.1 List of Variable Names and Definitions

`CPP_mean_amp` – The average pre-response amplitude of the CPP in each trial

`RT_S1` – The response time for S1 targets in seconds

`RT_S2` – The response time for S2 targets in seconds

`congruency` – Whether the S1 target and distractor were congruent or incongruent

`orientation_S1` – The direction of higher contrast gratings in the S1 target stimulus (left or right)

`orientation_S2` – The direction of higher contrast gratings in the S2 target stimulus (left or right)

`evidence_strength` – The contrast difference between higher and lower contrast gratings for S2 targets (weak, medium or strong)

`ID` – The participant ID number (used for defining random intercepts and slopes)

#### 2.2 Regression Model Equations

Full model equations are listed below. Here, we follow the conventions of the lme4 package notation (Bates et al., 2015) when describing the structure of mixed effects regression models.

##### 2.2.1 Testing for an Effect of S1 Congruency on S1 CPP Pre-response Amplitudes

For analyses of S1 CPP pre-response amplitudes, we fit linear mixed effects regression models. We compared the following model including effects of S1 congruency:

```
CPP_mean_amp ~ congruency + S1_orientation + (1 + congruency | ID)
```

With a comparison model that did not include a fixed effect of congruency:

```
CPP_mean_amp ~ S1_orientation + (1 + congruency | ID)
```

##### 2.2.2 Testing for an Effect of S1 RT on S1 CPP Pre-response Amplitudes

For analyses of S1 CPP pre-response amplitudes, we fit linear mixed effects regression models. We compared the following model including effects of S1 RT:

```
CPP_mean_amp ~ congruency + S1_orientation + RT_S1 +
(1 + congruency | ID)
```

With a comparison model that did not include a fixed effect of S1 RT:

```
CPP_mean_amp ~ congruency + S1_orientation + (1 + congruency | ID)
```

##### 2.2.3 Testing for an Effect of S2 Evidence Strength on S2 CPP Pre-response Amplitudes

For analyses of S2 CPP pre-response amplitudes, we fit linear mixed effects regression models. We compared the following model including effects of S2 evidence strength:

```
CPP_mean_amp ~ S1_orientation * S2_orientation + congruency
+ evidence_strength + (1 | ID)
```

With a comparison model that did not include a fixed effect of S2 evidence strength:

```
CPP_mean_amp ~ S1_orientation * S2_orientation + congruency +
(1 | ID)
```

#### 2.2.4 Testing for an Effect of S1 Congruency on S2 CPP Pre-response Amplitudes

For analyses of S2 CPP pre-response amplitudes, we fit linear mixed effects regression models. We compared the following model including effects of S1 congruency:

```
CPP_mean_amp ~ S1_orientation * S2_orientation + evidence_strength
+ congruency + (1 | ID)
```

With a comparison model that did not include a fixed effect of congruency:

```
CPP_mean_amp ~ S1_orientation * S2_orientation + evidence_strength +
(1 | ID)
```

### 3. Numbers of epochs in each condition that were included in EEG analyses

**Supplementary Table S1.** *Summary statistics for numbers of epochs retained for analyses per participant, split by condition for S1 and S2 targets.*

| Comparison | Condition | Mean Epochs | Median Epochs | SD Epochs | Min Epochs | Max Epochs |
|---|---|---|---|---|---|---|
| S1 Congruent and Incongruent | Congruent | 213 | 213 | 14 | 175 | 237 |
| | Incongruent | 184 | 183 | 21 | 148 | 223 |
| S1 Binned By Congruency and RT Quantile | Congruent Fast | 70 | 70 | 5 | 58 | 78 |
| | Congruent Medium | 70 | 71 | 5 | 57 | 78 |
| | Congruent Slow | 72 | 72 | 5 | 60 | 81 |
| | Incongruent Fast | 61 | 60 | 7 | 49 | 74 |
| | Incongruent Medium | 61 | 61 | 7 | 49 | 73 |
| | Incongruent Slow | 62 | 62 | 7 | 50 | 76 |
| S2 Binned by S1 Congruency and S2 Evidence Strength | S1 Congruent S2 Weak | 62 | 64 | 9 | 34 | 76 |
| | S1 Congruent S2 Medium | 67 | 66 | 7 | 42 | 79 |
| | S1 Congruent S2 Strong | 69 | 69 | 6 | 52 | 79 |
| | S1 Incongruent S2 Weak | 54 | 54 | 9 | 35 | 71 |
| | S1 Incongruent S2 Medium | 58 | 57 | 9 | 39 | 78 |
| | S1 Incongruent S2 Strong | 58 | 57 | 8 | 42 | 75 |

### 4. Mean RTs for Correct and Error Responses to S1 and S2 Targets

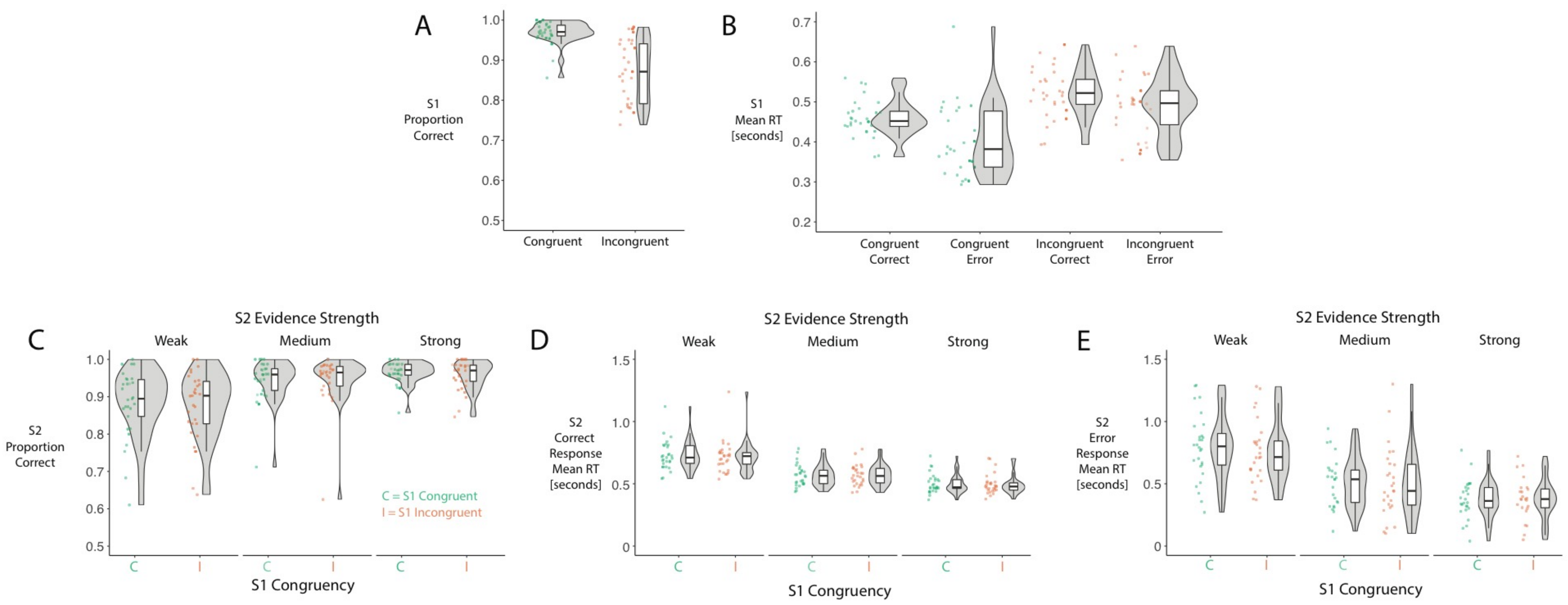


Supplementary Figure S1. Accuracy and mean RTs for correct and error responses following S1 and S2 targets. A) Proportions of correct responses following S1 targets with congruent and incongruent distractors. B) Mean RTs for correct and error responses to S1 targets with congruent and incongruent distractors. C) Proportions of correct responses to S2 targets, split by S2 evidence strength and S1 distractor congruency. D, E) Mean RTs for correct and error responses to S2 targets, split by S1 congruency and S2 evidence strength. Mean RTs for error responses within each condition were only calculated for participants who made at least one error in that condition.

### 5. Time-Frequency Plots for S1 Targets at Electrode FCz

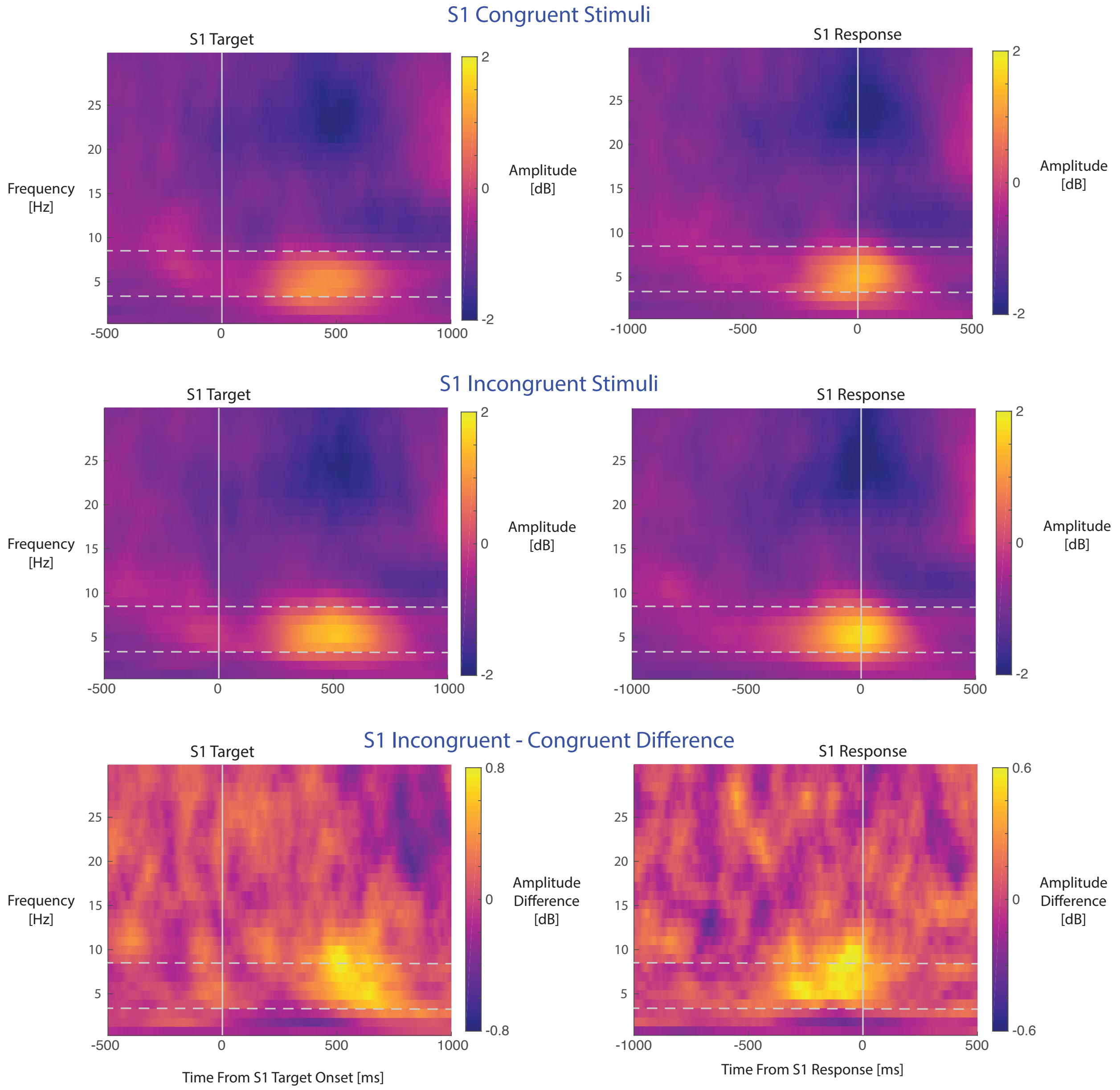


Supplementary Figure S2. Time-frequency plots relative to S1 target onset (left column plots) and the S1 response (right column plots) for data at electrode FCz, for congruent and incongruent S1 stimuli. Dashed horizontal grey lines denote the theta [4-8 Hz] band used for measurement of theta amplitude.

### 6. Lateralised [Contralateral – Ipsilateral] Mu/Beta Differences for S1 Targets

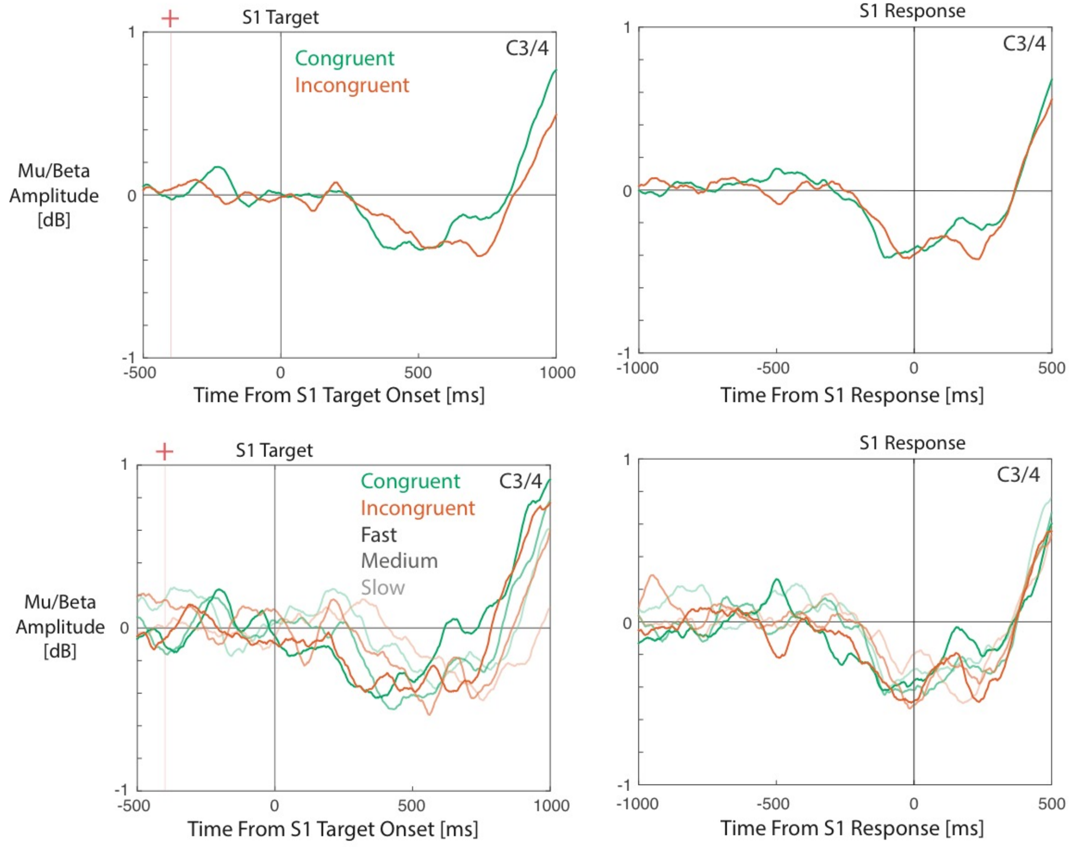


Supplementary Figure S3. Mu/Beta differences between electrodes contralateral and ipsilateral to the response hand, for correct responses to S1 targets. Mu/Beta amplitudes are plotted aligned to the S1 target onset (left panels) and to the S1 response time (right panels). Following S1 targets, contralateral and ipsilateral Mu/Beta amplitudes do not appear to diverge until approximately 200 ms preceding the time at which a keypress is registered.

### 7. Time-Frequency Plots for S2 Targets at Electrode FCz

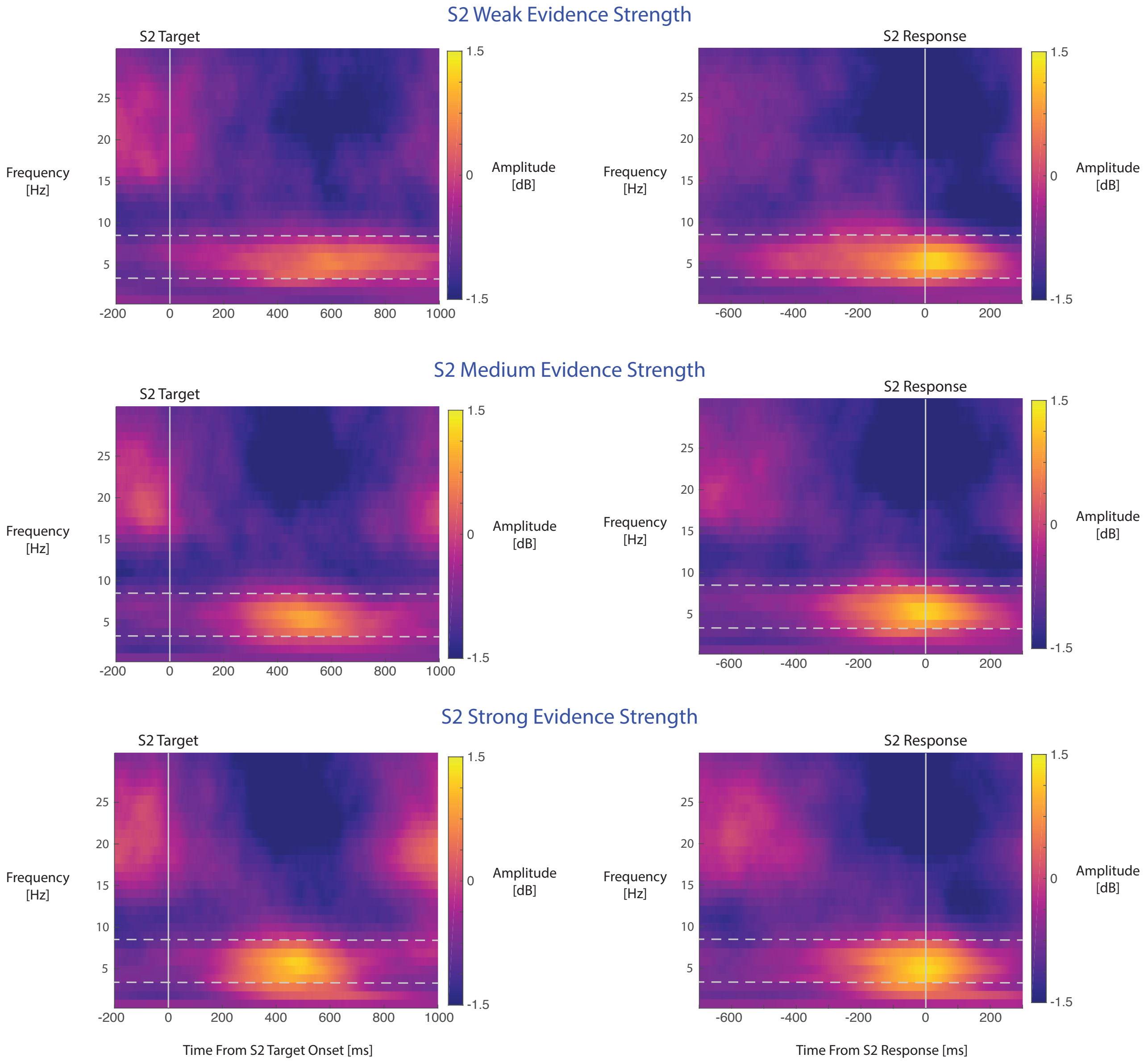


Supplementary Figure S4. Time-frequency plots relative to S2 target onset (left column plots) and the S2 response (right column plots) for data at electrode FCz, for weak, medium and strong evidence strength stimuli. Dashed horizontal grey lines denote the theta [4-8 Hz] band used for measurement of theta amplitude.

### 8. Supplementary Experiment 2 Methods

#### 8.1 Participants

29 participants (21 female, 8 male), aged between 18–37 (M = 22.8, SD = 4.2) took part in experiments 2A and 2B. Please note that Experiment 2 as reported in the main manuscript corresponds to Experiment 2B in this supplementary materials document. Participants were right-handed, fluent in spoken and written English, and had normal or corrected-to-normal vision. 4 participants were excluded from analyses for experiment 2A, and 1 participant was excluded from experiment 2B, because they achieved less than 60% accuracy in responding to one of the S1 or S2 target conditions. The order of participation in experiments 2A and 2B was counterbalanced across participants.

#### 8.2 Stimuli and Procedure

Stimuli, trial structure and hardware used for presenting stimuli and collecting responses were identical to Experiment 1, with the following changes. Experiment 2A included the same conditions as in Experiment 1, except that only stimuli of medium evidence strength (75/25% contrast levels) were presented as S2 stimuli (weak and strong evidence conditions were omitted). Therefore, there were two conditions: S1 congruent and S1 incongruent. This experiment was also shorter in duration than Experiment 1, and included 4 blocks of 48 trials each.

Experiment 2B was identical to Experiment 2A, except that an extra condition was added whereby the S2 stimulus was incongruent. Therefore, there were 4 conditions, corresponding to each combination of S1 and S2 congruent/incongruent status. There were equal numbers of trials for each condition within each block of the experiment. Experiment 2B included 6 blocks of 48 trials.

#### 8.3 Analyses of Behavioural Responses

Code used for analysing behavioural data in Experiment 2 will be made available at https://osf.io/eucqf/ at the time of publication. Accuracy and RTs were analysed in the same way as in Experiment 1 via comparisons of generalised linear mixed effects regression models. Fixed effects for analyses of S1 responses included S1 orientation and S1 congruency. Fixed effects for analyses of S2 responses included S1 congruency, and additive and interactive effects of S1 and S2 target orientation. The effect of S2 congruency, as well as the interaction between S1 and S2 congruency, were also included in analyses of data from Experiment 2B. The random effects structures were identical to models used in Experiment 1, including random slopes of S1 congruency and S1 target orientation for analyses of S1 responses, and a random intercept of participant for all S1 and S2 analyses.

As we found statistically significant interaction effects of S1 and S2 congruency on accuracy and RTs in Experiment 2B, we conducted follow-up model comparisons to investigate these interactions, using data from either S2 congruent or S2 incongruent trials in separate analyses.

### 9. Supplementary Experiment 2 Results

#### 9.1 Accuracy and RTs for S1 Targets

S1 target accuracy (i.e. proportion correct) scores and RTs are summarised in Figures S2 and S3. Models fit to data from experiments 2A and 2B that included the effect of S1 congruency provided better fits to the data for measures of accuracy (likelihood ratio test p values both $< 0.001$) and RTs (Experiment 2A $p < 0.001$, Experiment 2B $p = 0.003$). As in the original EEG experiment, participants were faster and more accurate when responding to congruent S1 stimuli across both experiments (as depicted in Supplementary Figures S2 and S3). Delta plots for S1 stimuli (in Figures S2C and S3C) were similar to those observed in existing studies (e.g., Ulrich et al., 2015).

#### 9.2 Accuracy and RTs for S2 Targets

In Experiment 2A (results depicted in Supplementary Figure S5), we tested for effects of S1 congruency on responses to S2 targets. There was no evidence for an effect of S1 congruency on participants' accuracy ($p = 0.958$) or RTs ($p = 0.946$) when responding to S2 targets.

In experiment 2B (results depicted in Supplementary Figure S6), there was an effect of S2 congruency on accuracy (likelihood ratio test $p < 0.001$), but little evidence for a fixed effect of S1 congruency ($p = 0.883$). However, a model including an interaction between S1 and S2 congruency fit better than a comparison model with only additive terms ($p = 0.032$), indicating that the magnitude of the S2 congruent/incongruent difference did depend on the congruency of the S1 stimulus, as is typical of congruency sequence effects (Gratton et al., 1992; Duthoo et al., 2013). Follow-up model comparisons using only trials from S2 congruent and incongruent conditions in separate analyses revealed that the effect of S1 congruency on accuracy was not statistically significant for analyses of trials with S2 congruent ($p = 0.120$) or incongruent ($p = 0.167$) targets. The interaction appeared to be driven by a difference in the sign of the effects in S2 congruent and incongruent trials. Fixed effects estimates indicated slightly worse accuracy following S1 incongruent compared to congruent stimuli for S2 congruent targets, but slightly better accuracy for S2 incongruent targets, as depicted in Supplementary Figure S6D.

For modelling of RTs in Experiment 2B, model fits were improved by adding fixed effects of both S1 and S2 congruency (both p's $< 0.001$). Moreover, there was a statistically significant interaction between effects of S1 and S2 congruency ($p < 0.001$).

We followed up this interaction effect by testing for effects of S1 congruency using data from either S2 congruent or incongruent trials in separate analyses. For analyses of data from S2 congruent trials, there was no evidence of an effect of S1 congruency ($p = 0.620$). For analyses of data from S2 incongruent trials the congruency of S1 did affect RTs, with faster responses following S1 incongruent stimuli ($p < 0.001$), as is typical of congruency sequence effects. Overall, it appeared that effects of S1 congruency were only found for trials where the S2 stimulus was incongruent.

These effects on RTs to S2 incongruent stimuli can also be observed in the delta plots in Supplementary Figure 3F, which display RT differences across congruent and incongruent conditions for each RT quantile, arranged from fastest to slowest RTs. When the S1 stimulus was incongruent, the slope of the delta plot was shallower than when the S1 stimulus was congruent. This is what would be predicted if one assumes that post-conflict adjustments to decision processes reflect a change in the rate of accumulation of decision evidence associated with sensory information provided by the S2 distractor (see the results and discussion sections in the manuscript).

Overall, these results demonstrate that congruency sequence effects can indeed be found using our experimental design. However, effects on RTs were only found for incongruent, rather than congruent, S2 stimuli.

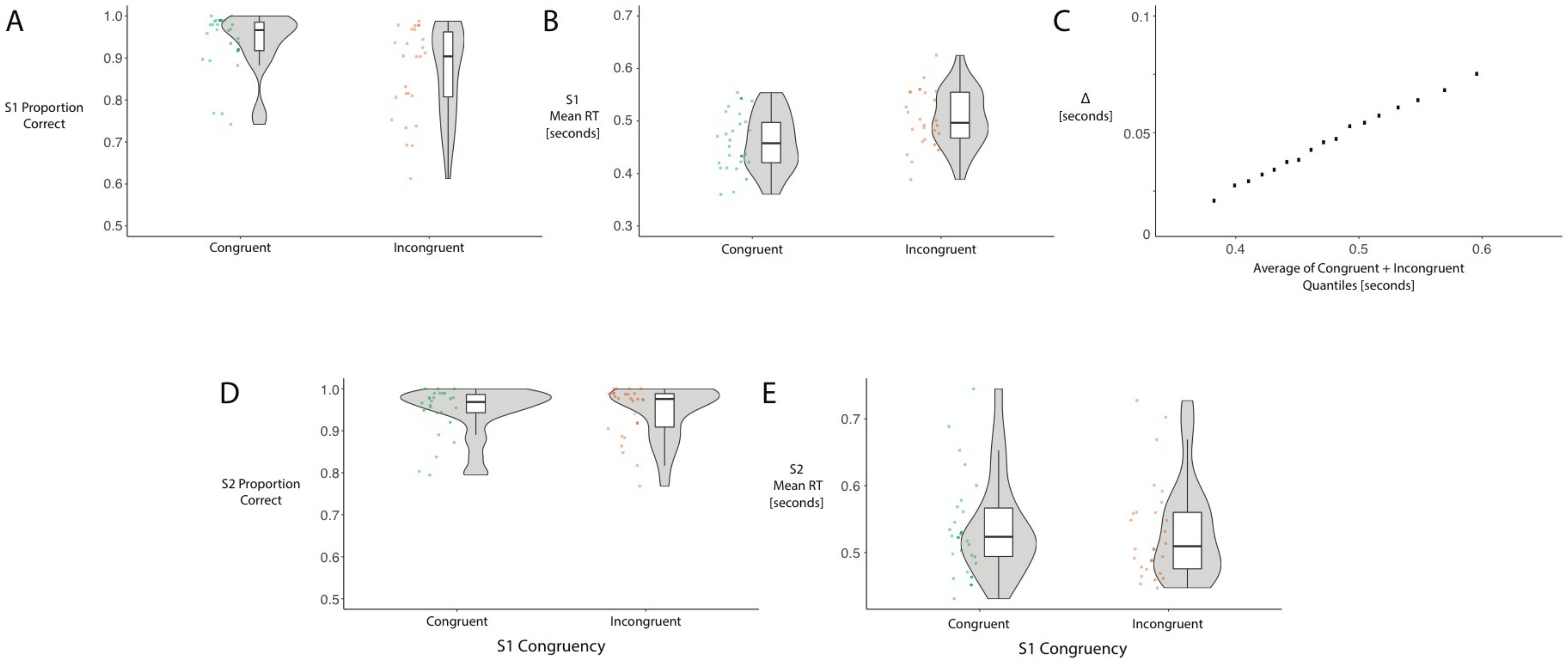


Supplementary Figure S5. Accuracy and mean RTs following S1 and S2 targets for experiment 2A. A) Proportions of correct responses following S1 targets with congruent and incongruent distractors. B) Mean RTs for correct responses to S1 targets with congruent and incongruent distractors. C) Delta plot displaying the [incongruent – congruent] differences in RTs for 10-90% quantiles in steps of 5% on the Y axis and the average of each congruent and incongruent condition RT quantile on the X axis. D) Proportions of correct responses to S2 targets split by S1 congruency. E) Mean RTs for correct responses to S2 targets, split by S1 congruency.

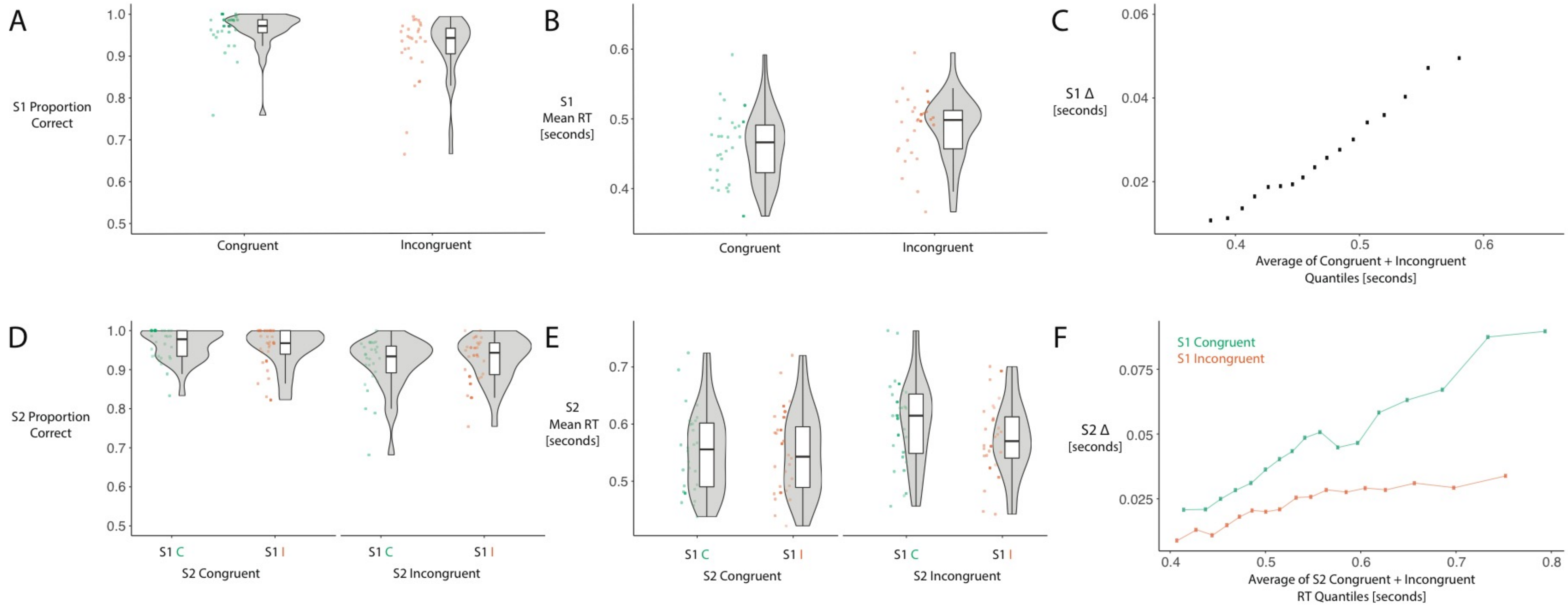


Supplementary Figure S6. Accuracy and mean RTs following S1 and S2 targets for experiment 2B. A) Proportions of correct responses following S1 targets with congruent and incongruent distractors. B) Mean RTs for correct responses to S1 targets with congruent and incongruent distractors. C) Delta plot displaying the [incongruent – congruent] differences in RTs for 10-90% quantiles in steps of 5% on the Y axis and the average of each congruent and incongruent condition RT quantile on the X axis. D) Proportions of correct responses to S2 targets split by S1 and S2 congruency. E) Mean RTs for correct responses to S2 targets, split by S1 and S2 congruency. F) Delta plots displaying the S2 [incongruent – congruent] differences for each RT quantile, plotted separately for S2 targets following S1 congruent and S1 incongruent stimuli.

### 10. Supplementary Material References


Bates, D., Maechler, M., Bolker, B., & Walker, S. (2015). Fitting linear mixed-effects models using lme4. *Journal of Statistical Software, 67*(1), 1-48.

Duthoo, W., Wurh, P., & Notebaert, W. (2013). The hot-hand fallacy in cognitive control: Repetition expectancy modulates the congruency sequence effect. *Psychonomic Bulletin and Review, 20*, 798-805.

Gratton, G., Coles, M.G.H., & Donchin, E. (1992). Optimising the use of information: Strategic control of activation of responses. *Journal of Experimental Psychology: General, 121*(4), 480-506.

Ulrich, R., Schroter, H., Leuthold, H., & Birngruber, T. (2015). Automatic and controlled stimulus processing in conflict tasks: Superimposed diffusion processes and delta functions. *Cognitive Psychology, 78*, 148-174.
